# Single cell expression and chromatin access of the *Toxoplasma gondii* lytic cycle identifies AP2XII-8 as an essential pivotal controller of a ribosome regulon

**DOI:** 10.1101/2023.10.06.561197

**Authors:** Jingjing Lou, Yasaman Rezvani, Argenis Arriojas, David Degras, Caroline D. Keroack, Manoj T. Duraisingh, Kourosh Zarringhalam, Marc-Jan Gubbels

**Author notes:** equal contribution / co-first authors (Jingjing Lou, Yasaman Rezvani). Department of Molecular Microbiology and Immunology, Brown University, Providence, RI, USA.

## Abstract

Sequential lytic cycles driven by cascading transcriptional waves underlie pathogenesis in the apicomplexan parasite *Toxoplasma gondii*. This parasite’s unique division by internal budding, short cell cycle, and jumbled up classically defined cell cycle stages have restrained in-depth transcriptional program analysis. Here, unbiased transcriptome and chromatin accessibility maps throughout the lytic cell cycle were established at the single cell level. Correlated pseudo-timeline assemblies of expression and chromatin profiles mapped transcriptional versus chromatin level transition points promoting the cell division cycle. Sequential clustering analysis identified putatively functionally related gene groups facilitating parasite division. Promoter DNA motif mapping revealed patterns of combinatorial regulation. Pseudo-time trajectory analysis revealed transcriptional bursts at different cell cycle points. The dominant burst in G1 was driven by transcription factor AP2XII-8, which engages TGCATGCG/A and TATAAGCCG motifs, and promoted the expression of a regulon encoding 40 ribosomal proteins. Overall, the study provides integrated, multi-level insights into apicomplexan transcriptional regulation.

## Introduction

The beginning and end-point of the *Toxoplasma gondii* lytic cycle is an invasion competent tachyzoite harboring the apicomplexan phylum-defining complex of apical secretory organelles and cytoskeletal structures [1]. Two daughter tachyzoites per cell division round are produced by internal budding (endodyogeny) [2]. In this process, two daughter buds encased by the cortical membrane cytoskeleton elements are assembled around the centrosome, which ultimately consumes the mother cell [3]. Apicomplexan cell division is strikingly different from mammalian cells and the canonical phases of the cell division cycle poorly fit this process. Notably, daughter budding, which can be considered cytokinesis, occurs simultaneously with S and M cell cycle phases (**Figure 1A**) [4]. A classically defined G2 phase is absent, whereas only G1 is clearly separated from all other events. To define apicomplexan cell division, we recently introduced the following cell division modules: 1. mother cytoskeleton disassembly, 2. DNA synthesis and chromosome segregation (D&S), 3. karyokinesis, and 4. daughter bud assembly (budding) [5]. Division modes within and across apicomplexan parasites differ by timing, recurrences of individual modules, and the number of repetitions, among which *T. gondii* endodyogeny is the simplest and most accessible model system.

**Figure 1.**
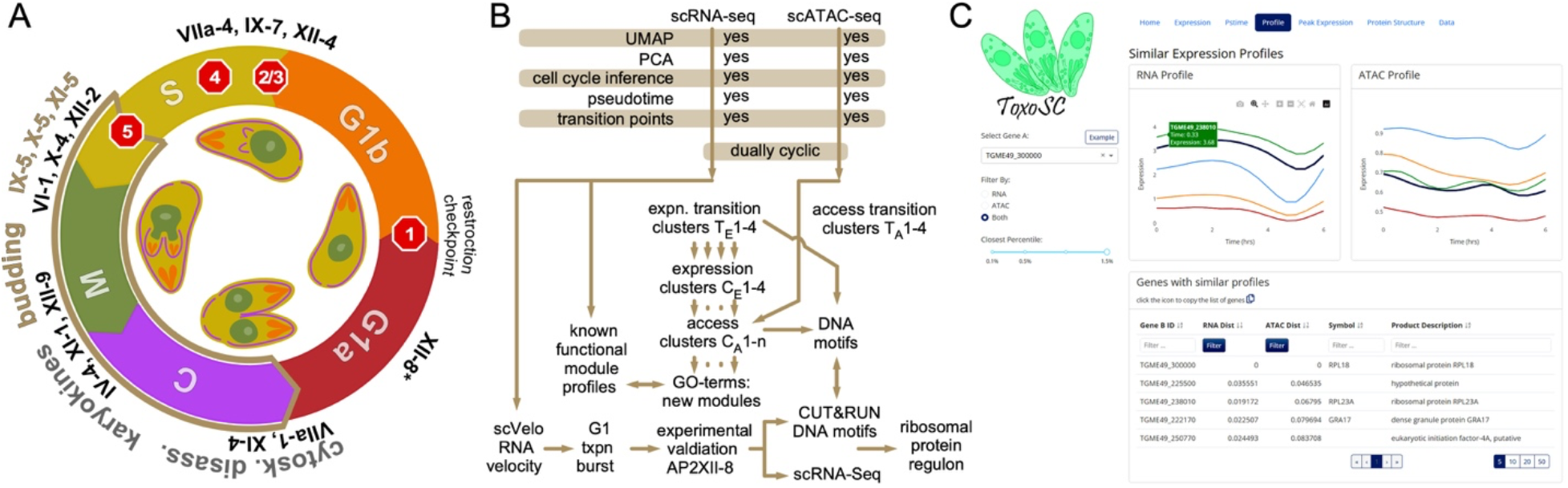
Overviews of the *T. gondii* tachyzoite lytic cycle and experimental approach. **A.** Schematic of the cell division cycle. Division by internal budding is illustrated in the inner circle and overlaps with S, M, and C phases. Stop signs illustrate the following checkpoints: 1. restriction; 2. DNA licensing; 3. centrosome duplication; 4. spindle assembly; 5. daughter assembly [42]. AP2 factors that have experimentally been validated are marked at the stages they act. * AP2XII-8 was established in this study. Panel modified from [16]. **B.** Schematic summary of the experimental and computational approaches taken in this study. **C.** Web-app displaying query for genes by similarity in expression and/or chromatin access pattern.

Cascading waves of ‘just-in-time’ gene expression profiles promote progression through the division cycle [2, 6]. The chromatin state defines the developmental stage and poises expression of the required genes throughout the cycle [2, 5, 7]. Temporal gene expression throughout the cell division cycle is mediated by transcription factors (TFs). Cell division modules comprised of clusters of co-regulated genes are controlled by several TFs and post-translational modifiers (PTMs). Each TF controls a ‘regulon’, defined as a set of genes under the regulation of the TF. The most abundant TFs in *T. gondii* belong to the apetala 2 (AP2) family, which is the key TF family driving the apicomplexan transcriptional waves [8]. Archetypical apicomplexan AP2 factors (ApiAP2s) are defined by 1-2 AP2 DNA binding domains and harbor no other functional domains [9–12]. AP2s are capable of homo- or hetero-dimerization and can act cooperatively or competitively as repressors and/or activators on the same gene [13–16]. *T. gondii* harbors 67 AP2 encoding genes [17–19].

Several studies have analyzed the transcriptome and chromatin state of apicomplexan parasites. For example, transcriptome oscillations throughout the lytic cycle of the *T. gondii* tachyzoite life stage have been determined by microarrays on synchronized parasites [18], and later by single cell RNA sequencing (scRNA-seq) on a small number of FACS sorted tachyzoites [6]. Moreover, bulk genome-wide studies on asynchronously cycling parasites have revealed that active promotor regions are marked by a complex pattern of histones H3 and H4 methylation and acetylation [20]. Together with recently applied ‘Assay for Transposase-Accessible Chromatin’ (ATAC) on bulk replicating tachyzoites [21] this demonstrated that chromatin opening facilitates TF access and is a major regulatory mechanism underlying gene expression. Overall, genome-wide studies have provided fundamental insights into molecular mechanisms underlying the progression of the apicomplexan cell division cycle. However, there is no time-resolved, simultaneous expression and chromatin access data available that permits the analysis of how different levels of control are integrated.

Here, we expand on the transcriptional insights by combining our recently developed experimental tools and computational models [22] with scRNA-seq data and single cell ATAC-seq (scATAC-seq) data on replicating tachyzoites (**Figure 1B**), generated in parallel. Using a custom pseudo-time analysis pipeline, we constructed the oscillating transcriptome and chromatin landscape of *T. gondii* and showed that expression and chromatin accessibility are highly correlated. Through analysis of gene expression patterns and open chromatin regions, we identified major transition points in the *T. gondii* cell division cycle, which revealed different organization levels of cell cycle checkpoint and progression regulations. Analysis of the cell cycle regulated AP2 TF expression profiles resolved four major clusters, among which, a cluster of AP2s peaks during the C/G1a transition was conjectured to fuel a major RNA explosion during G1a. Functional analysis of AP2XII-8, an essential TF driving this burst, revealed its requirement for the expression of a regulon of ribosomal genes. This was mediated by two DNA motifs contained within larger ribosomal protein motifs, indicating that AP2XII-8 cooperates with other factors to regulate ribosome gene expression. Data sets generated in this work provide a major resource to interrogate changes in gene expression program during *T. gondii* tachyzoite replication To facilitate usage and access to our datasets and tools, an interactive web-app is provided (hosted at https://umbibio.math.umb.edu/toxosc/) where users can interact with the data and perform various analyses, including clustering and co-expression analysis (**Figure 1C**).

## Results

### 1. Mapping *T. gondii* cell cycle progression by single cell RNA and ATAC sequencing

We performed 10x Genomics scRNA-seq and scATAC-seq on RH strain *T. gondii* tachyzoites throughout the lytic cell division cycle. For scRNA-seq we acquired 12,735 cells with an average of 6,220 reads/cell mapping to 488 genes/cell (median). scATAC-seq captured 3,506 cells at a depth of 68,670 reads per cell with a median of 2,911 fragments per cell (**Table S1**). The Seurat R package [23] was utilized to process and scale the data and perform dimensionality reduction. **Figure 2A-D** show the Uniform Manifold Approximation and Projection (UMAP) and principal component analysis (PCA) plots of the scRNA-seq and scATAC-seq data. We used the previously inferred canonical cell cycle stage assignments (G1a, G1b, S, M, C) based on DNA content of the 800 RH strain tachyzoites that were FACS sorted prior to scRNA-seq [6]. The distinct G1a and G1b phases are consistent with assignments based on older microarray data [18]. We transferred the inferred cell cycle phase labels to our data using canonical correlation analysis through the Seurat R package [23]. **Figure 2A** and **2C** show that the cell cycle progresses through a closed circular trajectory, sequentially traversing each cell cycle phase. This circular pattern is due to the periodic gene expression during the replication cycle.

**Figure 2.**
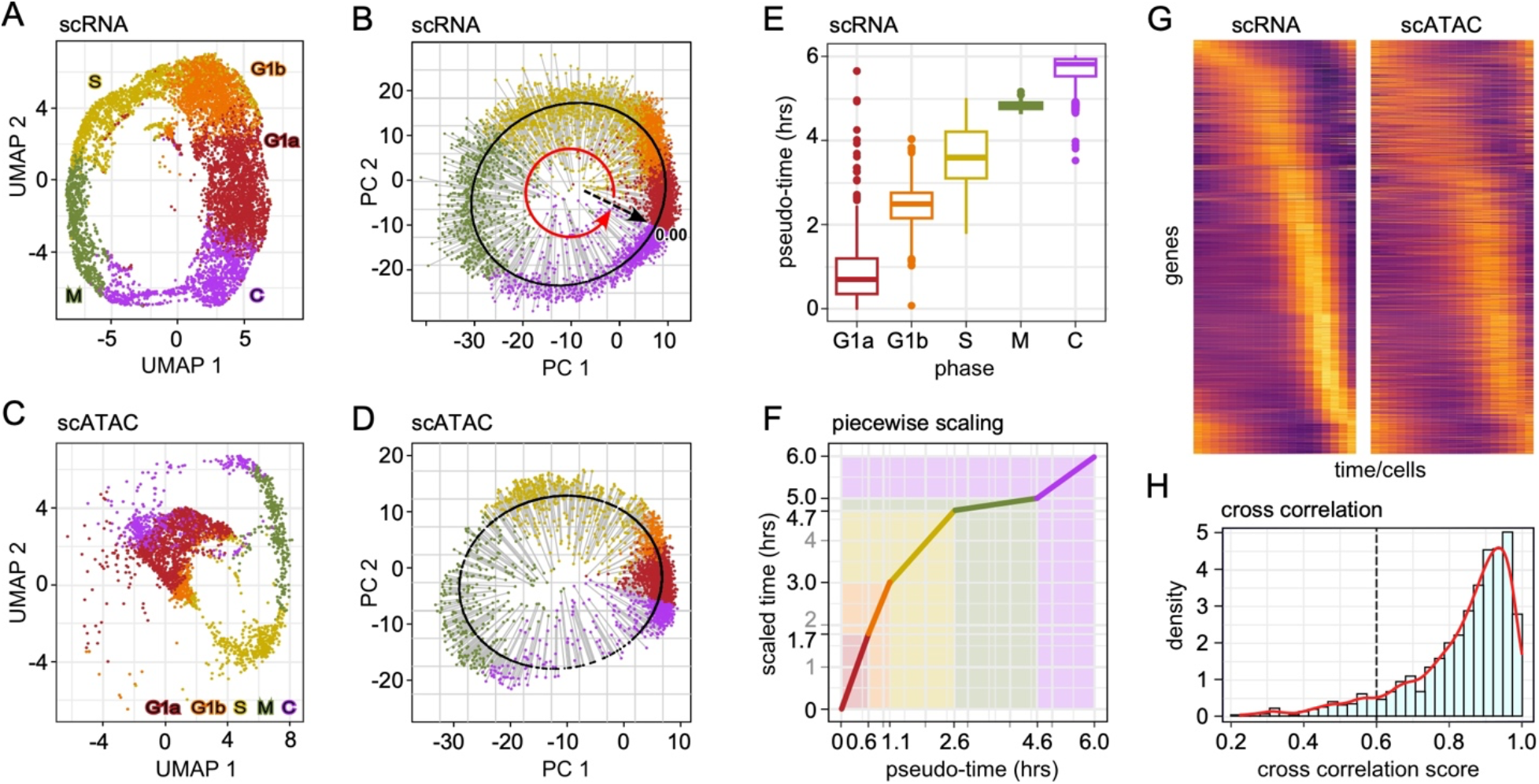
scRNA-seq and scATAC-seq analysis of the tachyzoite lytic cell division cycle. **A-D.** Asynchronously replicating intracellular RH strain tachyzoites were subjected to the 10x Genomics scRNA-seq **(A-B)** or scATAC-seq **(C-D).** Data was projected on UMAP **(A, C)** and the first two PCA **(B, D)** coordinates. Each dot represents a single tachyzoite. Derived cell division stage assignments were based on the definition put forward by Xue *et al* [6]. In panels **B** and **D**, the lytic cycle pseudo-time was reconstructed. The pseudo-time gene curve was constructed by mapping the cells ordered to 0–6 hrs, starting at the derived C-G1a transition. **E.** Box-plot representing distributions of cells at respective pseudo-time phases. Whisker indicated min/max and dots represent outliers. **F.** Calibration of scRNA-seq derived pseudo-time against experimentally defined timeline of the cell division cycle stages (scaled time) [4, 6]. **G.** Pseudo-time heatmaps of single cell transcriptome (left) and chromatin access (right) for the 1,238 genes displaying cyclic expression profiles. Rows: individual genes; columns: time (20 arbitrary time points) measured in different cells. Top to bottom organized by timing of peak gene expression, corresponding with the gene order in scRNA-seq. See **Figure S1C** for the reciprocal alignment by scATAC-seq peaks. **H.** Distribution of cross-correlation scores (CSS) representing the similarity of expression and accessibility curves across 1238 cyclic DEGs. The majority of genes show a CCS between their expression and accessibility profiles (CCS > 0.6). See also **Figure S1** and **Table S1**.

To map the progression of the cell cycle we performed a pseudo-time analysis by fitting a closed principal curve to the first two principal components (PCs) and orthogonally projected the cells onto this curve (**Figure 2B** and **2D**). The start of the cell cycle was set to the beginning of G1a, and cells were ordered along the trajectory within 20 evenly distributed time points. The arc-length was scaled from 0 to 6, reflecting the hours of the cell division cycle [4] (**Figure 2E**). We then applied a piecewise linear scaling to match the length of each pseudo-time phase to previously reported time-length of each phase [4] (**Figure 2F**). Subsequently, we generated expression and accessibility time-series curves for each expressed gene (∼6600 out of ∼8200 total detected genes) using the scaled pseudo-time by fitting weighted periodic smoothing splines to the expression and accessibility data. A Fourier-based analysis was used to identify cyclic genes, defined as those with significant oscillation determined by the magnitude of the dominant frequencies (i.e., the frequency with the highest amplitude) over one cycle. The analysis identified 4,097 cyclic genes according to expression curves and 3,506 genes with cyclic chromatin accessibility, of which 2,549 genes were dually cyclic in expression and accessibility curves (**Figure S1A**). Among the 2,549 dually cyclic genes, we identified genes significantly over expressed in at least one of the canonical phases (FC > 1.5, adj-p-value < 0.05), resulting in 1,238 unique cyclic differentially expressed genes (DEGs), out of 1,620 total detected DEGs (**Figure S1B, Table S2**). Taken together, we have assembled a unique data set of DEGs for which both expression and chromatin access patterns are available permitting correlative analysis of these two major drivers of *T. gondii* cell cycle progression.

### 2. Transition points of accessibility and expression, and charting regulatory events

Considering the just-in-time gene expression principle in Apicomplexa, the peak of expression of cyclic marker genes strongly correlates with when the gene product is needed in each step of the cell division cycle process, whereas the peak of chromatin access should correlate with the transcriptional and/or epigenetic switches in cell division cycle progression. To determine how these levels of control contribute to each cell cycle stage, we calculated the peak expression and accessibility times for each of the 1,238 cyclic DEGs. **Figure 2G** shows a heatmap of gene expression and chromatin accessibility ordered by peak expression time interval for the cyclic DEGs. Genes are expressed/become accessible sequentially with cascading peaks that span the entire cell division cycle. Accessibilities of genes show a similar pattern to expression (**Figure S1C**). To quantify this, we calculated that the curve cross-correlation score (CCS) between expression and accessibility curves. The CCS is above the 0.6 significance cut-off for over 90% of the dually cyclic genes (**Figure 2H)**.

We observed that peak expression times do not progress linearly (**Figure 2G**), indicating that there are points of inflection where a shift occurs in peak expression of gene groups. These points can define functional modules and switches in the cell division cycle. To pinpoint these transitions, we calculated the sequential peak times of cyclic DEGs (**Figure 3A** top) and identified the points of inflection (**Figure 3A** bottom). We utilized the points of inflection as ‘transition’ times, allowing us to group genes into four distinct expression clusters: T_E_1-4 (**Figure 3B**). Similar analysis was performed for peak accessibility, which established four chromatin access clusters named T_A_1-4 (**Figure 3C, D**). When overlaid with the inferred cell cycle annotations [6] (**Figure 3E** inner circle), we observed that expression transitions corresponded well with transitions from G1b to S and from S to M, but the transition from M to C was offset (**Figure 3E**). The T_E_1 cluster encompasses the entire G1a and G1b phases, whereas the ATAC-derived T_A_1 to T_A_2 transition marks the G1a to G1b switch, indicating regulation at chromatin level. The T_E_2 phase almost entirely matches the S phase, while the T_E_3 phase starts at the M phase and continues until mid C. The expression level regulation in these stages is more granular than the chromatin access cluster T_A_2, which encompasses G1b and S to the budding onset checkpoint (#5 **Figure 1A**). Cluster T_A_3 starts slightly prior to T_E_3 and ends roughly at the same spot. Finally, the T_E_4 and T_A_4 span from mid C back to early G1a with peak expression of clusters close to the transition boundary.

**Figure 3.**
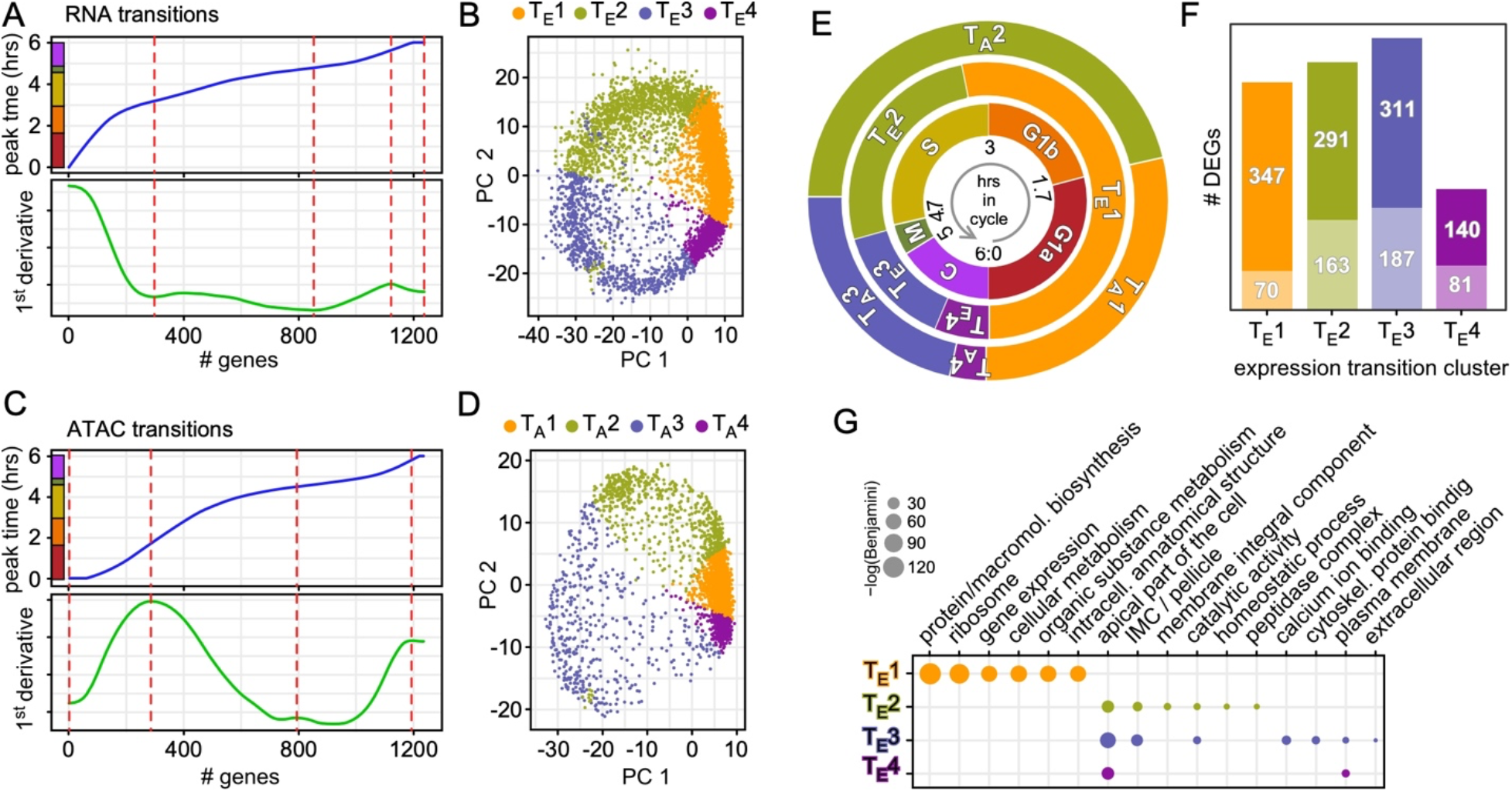
Transition points mapping in scRNA-seq and scATAC-seq data. **A, C.** Transcriptome **(A)** or chromatin access **(C)** transitions identified from peak expression or chromatin accessibility of 1,238 cyclic genes. Top panel shows the peak expression or chromatin accessibility time point for each gene ordered as in Figure 2G or **Figure S1C**; bottom panel shows the first derivative. Red-dotted lines mark the transition points. **B, D.** RNA or ATAC transitions mapped to PCA of scRNA-seq or scATAC-seq data, respectively. T_E_1-4 and T_A_1-4 mark the cell populations falling with each transition cluster. **E.** Representative illustration of the biologically assigned stages (inner circle), the expression level transitions (T_E_1-4) (middle circle), and the chromatin accessibility transitions (T_A_1-4) (outer circle). **F.** DEG analysis of each expression transition clusters T_E_1-4 defined in panels **A, B,** relative to the other clusters. The total number of genes is composed of the upper number representing the number of genes with manually enhanced annotation on ToxoDB and the lower number in the lighter shade representing the genes annotated as ‘hypothetical’. **G.** Functional enrichment for T_A_1-4 was performed by gene ontology (GO) analysis using ToxoDB annotations. Significant GO terms are shown (Benjamini < 0.1). See also **Table S2**.

Considering the cascading transcriptional events driving *T. gondii*’s replication, we theorized that the cell cycle switches were predominantly a reflection on expression transitions, with each transition propelled by a unique functional group of genes needed in each particular cell division step. To this end, we performed a differential gene expression analysis (DGEA) on the four expression derived clusters, which identified 1,347 unique DEGs (out of 1,590 total DEGs: **Figure 3F, Table S2**), followed by gene ontology (GO) term analysis. This revealed that T_E_1 contained genes typically associated with G1, whereas T_E_2-4 resolved in different functional clusters (**Figure 3G**). Although most GO terms did inform directly on specific functions, we did resolve two GO terms associated with daughter budding: ‘apical part of the cell’ and ‘IMC/pellicle’ spanned T_E_2-4, reflecting budding throughout S-M-C. Apicomplexan specific biology is also reflected in the fact that only 17% of the (G1) genes in T_E_1 are annotated as hypothetical, but that 35-38% of the T_E_2-4 genes are hypothetical since they comprise the defining apicomplexan specific organelles and processes.

### 3. Functional dissection by process and structure

We assembled lists of genes acting in the same process and/or structure to probe our data for modules comprising functionally related gene sets. We selected processes and structures within the overlapping S-M-C and budding events (**Figure 4A-C, Figure S2**). We did detect multiple expression profiles for most gene groups using automated clustering, though their peaks are typically around the same time point (**Figure 4C, Figure S2**). However, for a few gene sets we observed fairly tight expression patterns contrasted with considerable variation in chromatin access. We selected two representative gene sets for each of the three variations (loose, tight and multi-functional gene groups) between expression and chromatin access (**Figure 4A, B**). S- and M-phase genes represent ‘loose’ correlation. S-phase genes show tightly coordinated expression, peaking right around 3 hrs, which is the established onset of S-phase (**Figure 4A, C**). Expression of the M-phase, kinetochore protein encoding genes [24, 25], peaks just behind the S-phase genes (3-4 hrs; **Figure 4A**, **Figure S2A**). On the other hand, the chromatin access profile peaks for these S and M genes do not cluster well, with some peaking even in early G1 (**Figure 4A**). When the whole curve shape is considered (CCS), several genes in each group do drop below the 0.6 cut-off, confirming this ‘loose’ pattern (**Figure 4B**). Overall, loose chromatin access coordination despite a tight expression pattern suggests temporal control of TF activity, i.e., TF presence or post-translational modification.

**Figure 4.**
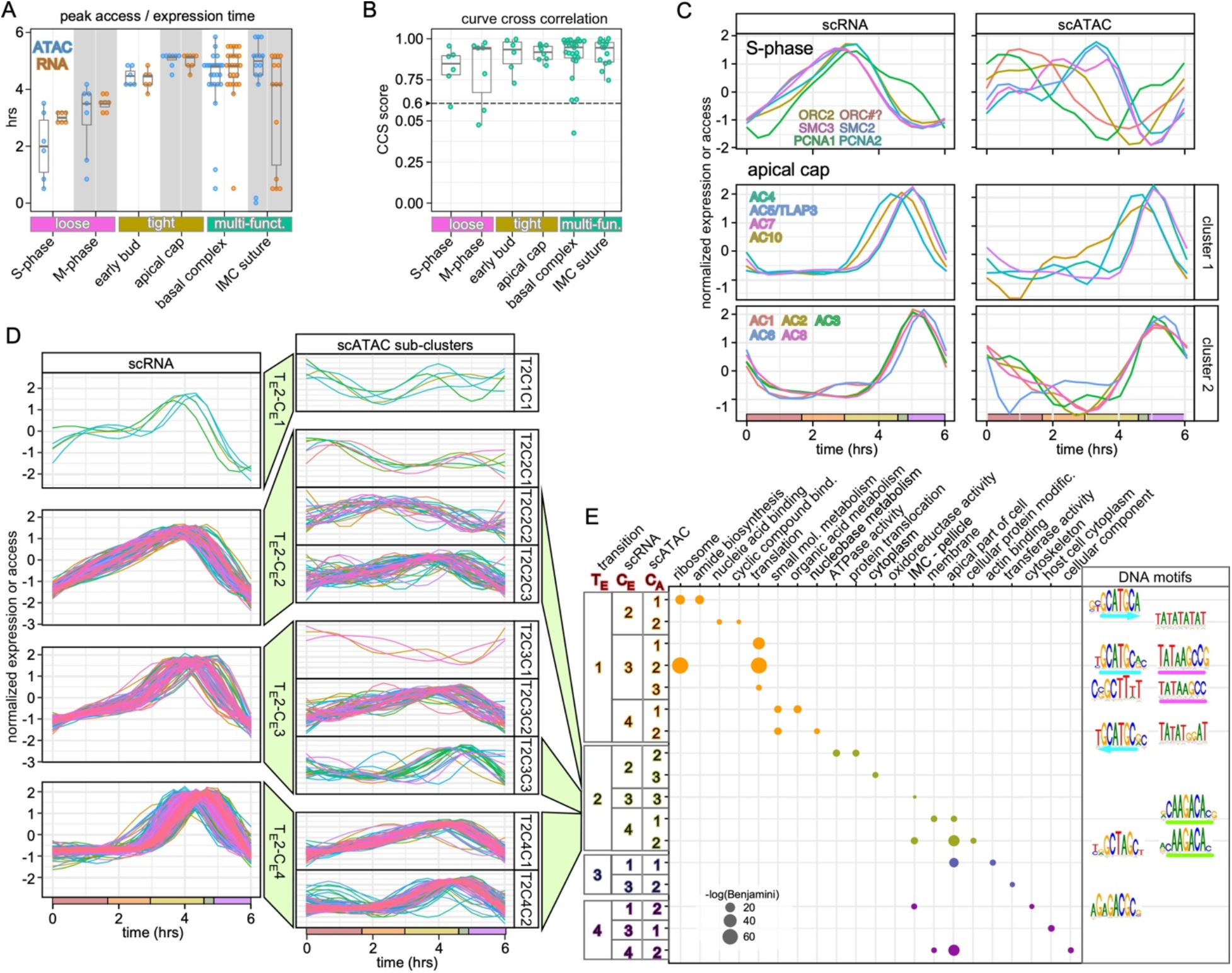
Dissection of functional gene sets expression and chromatin correlation. **A, B.** Selected functional gene sets (details in **Figure S2** and **S4**) representative of the different pattern (’loose’, ‘tight’, ‘multi-fun(ct).’) observed for peak times **(A)** and curve correlations between chromatin access and expression profiles **(B)**. Multi-fun(ct).: gene multi-functional sets in the same structure whose composition changes progressively through cell division. CCS > 0.6 indicates significant correlation. **C.** Representative S-phase and apical cap gene sets, displaying loose and tight coordinated expression and chromatin patterns, respectively. S-phase: origin recognition complex (ORC2 and ORC#?) [5], structural maintenance of chromosome factors SMC2 (condensin) and SMC3 (cohesin) [54], and proliferating cell nuclear antigens 1 and 2 (PCNA1 and PCNA2) [55]. **D.** Clustering analyses of DEGs in the T_E_2 transition cluster from Figure 3F. For complete analysis results, see **Figure S4**. **E.** GO term analysis using genes within each ATAC sub-cluster shown in Figure 4. Specific DNA motifs identified in the ATAC regions for genes in each sub-ATAC cluster are plotted. See also **Figure S3-5** and **Table S3**.

For the ‘tight’ correlation we selected genes encoding early daughter bud (**Figure 4A, B, Figure S2B**) and apical cap (**Figure 4A-C**) proteins, which are both very defined cytoskeletal elements assembled early in daughter budding. The apical cap genes show tight peak times (4.5-5 hrs) that correspond at the expression and chromatin level (**Figure 4C**). Thus, this tight correlation suggests coordinated chromatin and expression controls for these genes.

The ‘multi-functional’ gene sets represent the basal complex (BC) and different aspects of the IMC skeleton. The BC has several sequential functions through the lytic cycle, which is accompanied by a sequential change in composition [26, 27]. Indeed, we see this reflected in the relatively large window of peak times (**Figure 4A, Figure S2C**). A similar sequential assembly applies to the alveolin-domain IMC proteins [28] (**Figure S2D**). Despite an outlier, we do see good curve cross correlation for these gene sets. An additional gene set that our analysis surprisingly put in this group is the IMC suture proteins, which are assembled during daughter budding [29, 30]. Here we see a striking variation in expression peaks, whereas the curve cross-correlations are tight (**Figure 4A, B, Figure S2G**). Thus, this suggests the suture genes can be assigned to distinct expression clusters, most likely reflecting distinct biological roles. Taken together, our analysis provided a basis for future experimental work to dissect these predicted distinct functions.

Since transcriptional waves are AP2 driven, we examined the expression and chromatin access of the 32 cyclically expressed AP2s (**Figure S3**). They resolve in four expression clusters and largely match our recent analysis of the microarray time-course gene-expression data matching known functions of experimentally probed AP2 factors [16, 18]. In general, expression profiles are well mirrored in the chromatin access profiles, but there are some exceptions, e.g., AP2XI-3 and AP2VIIb−3. This dichotomy between the different co-ordination profiles as seen in a minority of cell division genes is also reflected in the TFs. Thus, although in general over 90% of cyclic genes show a cross correlation of expression and chromatin profiles (**Figure 2H**), some functional gene clusters deviate from this rule, which is important to keep in mind when interpreting automatically clustered data, and hints at more complex expression control mechanisms.

### 4. Lytic cycle dissection by transcription and chromatin patterns

We agnostically dissected the single cell data sets to gain further biological insights in coordinated gene clusters. We resolved the gene expression transitions groups of the DEGs (1,347 total based on expression-derived transition points, representing 33% of all 4,097 cyclically expressed genes) by an automatic time-series clustering (**Table S3**). Within each transition phase (T_E_1-4), the expression profiles of DEGs were further sorted into four distinct groups using a shape-based kernel method (**Figure 4D** and **Figure S4, left panels**), with expression peaks in each scRNA transition sub-cluster progressively shifting through each transition phase. The corresponding chromatin accessibilities for genes in each sub-cluster were assembled (**Figure S4**, middle panels). Concurring with previous observations that chromatin access profiles were more divergent than the expression levels, we found an indispensable need to perform a second clustering analysis, resulting in scATAC sub-clusters (right panels **Figure 4D** and **Figure S4**). This revealed a finer grain resolution within the transcriptional clusters. Thus, our data resolves at sequential levels: 1. scRNA transition; 2. Expression sub-clusters; 3. Chromatin sub-clusters. Biologically, this fits with the overlap of functional modules within the *T. gondii* cell division process which can be resolved in our data.

To test this model, we performed a cluster-based GO term analysis to assess the functions of clusters within each ATAC sub-phase (**Figure 4E, Table S3**). Results indicate cascading and partially overlapping processes. This analysis is consistent with the overlapping cell division processes defined by concurrently expressed functional modules that we can resolve in our data sets. To permit data queries with any gene of interest, we created a web app facilitating searches and clustering based on curve shape, either on the transcription or access level, as well as in combination (**Figure 1C**).

To add yet another dimension to the above assessments and inferences, we mapped conserved DNA motifs under each ATAC sub-cluster peak (**Figure 4E**). In parallel, we also performed motif search on the entire gene sets in the entire T_E_1-4 transition clusters to capture higher levels of co-accessibility (**Figure S5A**). This identified multiple motifs with high confidence, indicating that co-expressed/co-accessible genes are under the same regulatory controls at the DNA level. Strikingly, very similar motifs were found in distinct clusters. For example. the core motif GCATGC is found in clusters T_E_1C_E_2C_A_1, T_E_1C_E_3C_A_2, and T_E_1C_E_4C_A_2 (**Figure 4E**). Although these 3 clusters share this motif, each is unique in that it is either the only motif (T_E_1C_E_2C_A_1), or there is a combination with another unique motif (TATAAGCCG in T_E_1C_E_3C_A_2 and TATATGGAT in T_E_1C_E_4C_A_2), offering the fine-tuned regulation of distinct gene sets. The GO-terms associated with each of these three clusters do in part overlap between T_E_1C_E_2C_A_1 and T_E_1C_E_3C_A_2, which share the ‘ribosome’ GO term, but the GO-terms associated with T_E_1C_E_4C_A_2 revolve around nucleotide metabolism. Taken together, combinatorial DNA motifs permit fine tuning the expression of different functionally related gene sets.

Several of the other identified motifs mapped to previously identified AP2 DNA binding sites. Motif ‘GCTAGC’ (T_E_2-motif 5 and T_E_3-motif 8) as well as motif ‘CAAGACA’ have been reported as binding sites of AP2XI-5 [31] (**Figure S5A**). AP2XI-5 cooperates with AP2X-5 and is known to regulate multiple S/M-phase genes [15, 31]. T_E_2-motif 6 ‘CC(C/T)CCCC(C/T)’ matches closely to the previously reported binding site of AP2IV-4 (‘CACCCCCC(C/T)’), which is exclusively expressed in *T. gondii*’s tachyzoites in late S phase and functions in the suppression of early bradyzoite formation [32]. Other motifs, including the TATA box motifs, have been described before but have no known role [21]. Furthermore, T_E_4-motif 10 (A/TGAGACG) is a sequence element found in dense granule (GRA) and SAG1 promoters [33]. Finally, we mapped the distance of the identified motifs from the transcription start sites (TSSs) to show that all motifs are within 2 kb around the TSSs (most even within 1 kb), strongly supporting the interpretation that these are *cis*-motifs acting on the nearest gene (**Figure S5B**).

### 5. AP2XII-8 is required for proper progression through the G1a phase

RNA velocity, derived from the kinetics of the ratio between spliced and incompletely spliced transcript, can be used as a measure of transcription rate [34, 35]. Bursts in transcription rates point at major switches or checkpoints in transcriptional profiles [35]. We subjected the scRNA-seq data set to scVelo analysis [35]. **Figure 5A** shows the magnitude of cell velocities and reveals a major peak in G1a, next to one or two modest bursts in S-phase and a sharper burst at the M-C transition.

**Figure 5.**
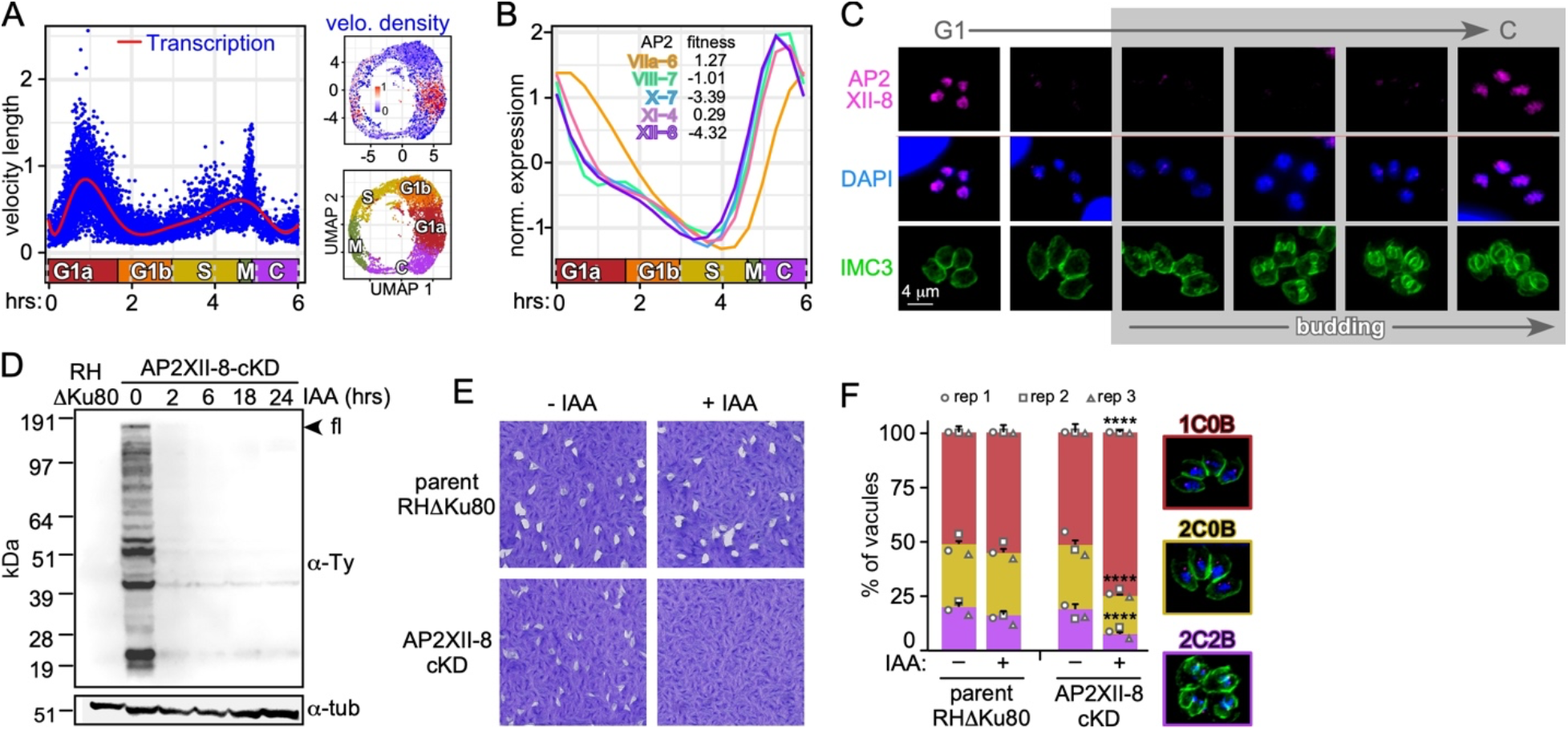
Functional characterization of AP2XII-8 demonstrates it is essential for G1 progression. **A.** scVelo plot depicting RNA velocity length along the pseudo time. The calculated RNA velocity length can be viewed as a measure of active transcription. Each dot represents velocity length of a cell. The red curve shows a fitted smoothing spline to the data. Inserts on the right show UMAPs of the density of RNA velocity (top) and the cell cycle stages as reference (bottom). **B.** C-G1 peak expression profiles of the cluster of 4 cyclically expressed AP2s (**Figure S3**). Fitness score for each of the five AP2s is listed. **C.** Protein expression and localization of endogenously 5xTy tagged AP2XII-8 (strategy and validation: **Figure S6**) throughout the lytic cycle by IFA. AP2XII-8 was visualized with Ty MAb, DAPI highlights DNA and IMC3 antiserum tracks the cytoskeleton of mother and daughter buds. **D.** AP2XII-8-mAID-5xTy knock-down kinetics assessed by western blot. Blot was probed with Ty MAb BB2 or α-tubulin as loading control. Full length (fl) AP2XII-8-mAID-5xTy has a predicted MW of 180.3 kDa. **E.** Plaque assay of AP2XII-8 depletion (8 days). Representative of 3 biological replicates is shown. **F.** Phenotype quantifications in 24 hr treated RHΔKu80 and AP2XII-8-cKD parasites. 1C0B: 1 centrosome, 0 buds (G1); 2C0B: 2 centrosomes, 0 buds (S); 2C2B: 2 centrosomes, 2 buds (M/C). Centrosomes were detected using α-Centrin. At least 100 vacuoles were counted per biological replicate. A total of 3 replicates per experiment. Statistics were performed using a z-test. **** *p* < 10^-4^. See also **Figure S6** and **Table S7**.

In particular the G1a RNA velocity burst intrigued us as this stage has not been the focus of studies in Apicomplexa. We reasoned that this burst must be controlled by cyclic AP2s whose expression precedes G1a. AP2 cluster 4 comprised five AP2 TFs with such profiles (**Figure S3**). Two of these AP2 factors have a tachyzoite fitness score below -3, suggesting they are essential for lytic cycle progression [36]: AP2X-7 and AP2XII-8 (**Figure 5B**). We had previously found that AP2XII-8 expression is enriched in a G1 entry temperature sensitive mutant [37]. Therefore, we selected AP2XII-8 to probe into the G1 transcription by experimentally dissecting AP2XII-8 function.

AP2XII-8 RNA expression peaks at the C-G1 transition (T_E_4-T_E_1) (**Figure 3E, 5B**). We tagged the endogenous AP2XII-8 by CRISPR/Cas9 mediated insertion of a mini auxin-inducible degron (mAID) tag in frame with 5xTy epitopes (**Figure S6A**). We tracked AP2XII-8 expression through the cell division cycle by immunofluorescence assay (IFA) co-stained with IMC3 as a daughter budding marker and DAPI as reporter for S-phase progression (nucleus size). AP2XII-8 localizes to the nucleus, as expected from a TF, and its temporal profile mimics its RNA expression: only detectable during the C-G1 transition (**Figure 5C**). Next, we demonstrated that AP2XII-8 could be depleted from the parasites by adding IAA for as little as 2 hours (**Figure 5D**). We did not observe any proliferation by plaque assays which confirms that AP2XII-8 is essential for the lytic cycle (**Figure 5E**). Subsequently, AP2XII-8 depletion results in an accumulation of parasites with a single centrosome and no daughter buds, supporting a role for AP2XII-8 in G1 progression (**Figure 5F**). Thus, these findings pinpoint AP2XII-8 as an essential driver of the G1 transcriptional burst.

### 6. AP2XII-8 target identification by CUT&RUN and scRNA-seq

To resolve how AP2XII-8 promotes G1 progression we sought to determine the genes under its control. We applied <u>C</u>leavage <u>U</u>nder <u>T</u>argets and <u>R</u>elease <u>U</u>sing <u>N</u>uclease (CUT&RUN) to determine the sites in the genome occupied by AP2XII-8. Although Ty epitope mediated CUT&RUN has been applied before to *T. gondii* [38], we first optimized the procedure. We harvested two different numbers of parasites (200 and 500 million) and titered out two different antibodies. We determined that 200 million parasites with 2 μg BB2 antibody had the highest detection power (**Figure S7A-D**). We processed the CUT&RUN data using four negative controls and identified 2,338 binding events (peaks) with high stringency (adj_pvalue < 0.05). 88% of CUT&RUN and 94% of ATAC peaks fall within 2 kb distance of the TSS, consistent with a role in transcriptional regulation (**Figure 6A**). We merged the nearby CUT&RUN peaks and assigned the long peaks to the nearest downstream gene, creating a list of 970 genes targeted by AP2XII-8 (**Table 4**). The binding sites of 940 (97%) of these genes fall within the peaks determined from the scATAC-seq data (5,322 peaks; **Figure 6B, Table S5**). If we apply an even more stringent definition of direct targets, i.e., the peak is not detected in any of the negative controls’ 246 genes are found to be engaged by AP2XII-8, which is reflected in their 5-fold stronger read count intensity compared to the rest of the genes (**Figure S7E**). Robust overlaps between the CUT&RUN and the ATAC accessibility profiles can be seen in the representative chromosome maps of two different genes, where AP2XII-8 is detected in the promoter (**Figure 6C**).

**Figure 6.**
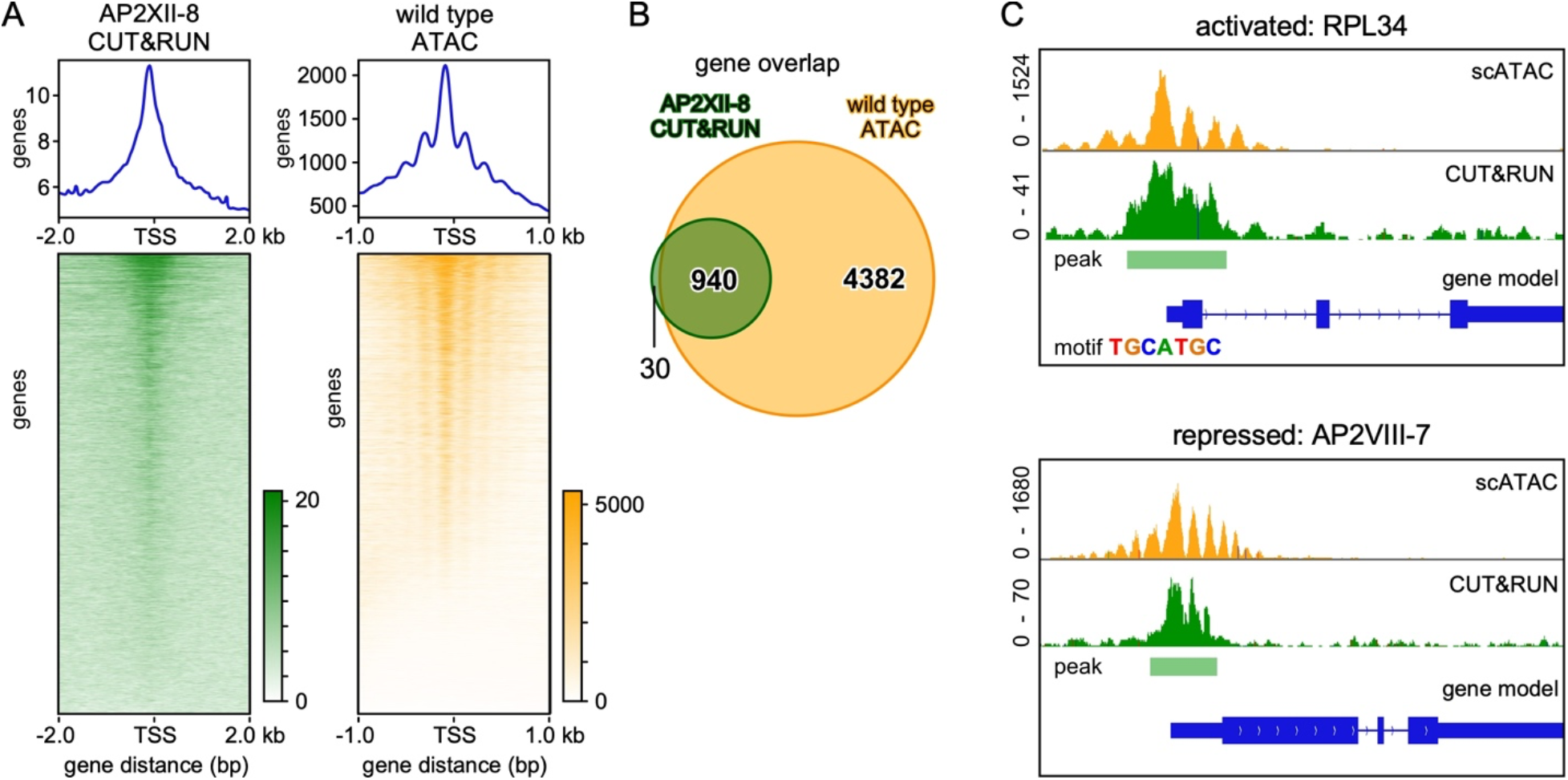
Identification of AP2XII-8 DNA binding motifs by CUT&RUN. **A.** Display of AP2XII-8 CUT&RUN (left) and wild type ATAC-peaks ranked by peak height (top to bottom) around the transcriptional start site (TSS). 2,338 AP2XII-8 CUT&RUN peaks were mapped upon concatenating the highly significant (adj_pvalue < 0.05) binding events using 4 data sets (AP2XII-8 vs 4 negative controls; **Figure S7E**). **B.** Venn diagram shows the overlap of 970 genes to which CUT&RUN peaks were assigned with the 5,322 genes with an ATAC profile from the scATAC-seq data considered as bulk (constitutive and cyclic at all time points combined) **C.** Two representative examples of genes with a CUT&RUN and ATAC profile around their TSS. RPL34 (TGGT1_227600) is activated by AP2XII-8 and contains a TGCATGC (motif 2) in its promoter (see Figure 7); AP2VIII-7 (TGGT1_269010) is suppressed by AP2XII-8 and is devoid of motif 1 or 2. See also **Figure S7** and **Tables S4, 5**.

Several processes mediate TF-DNA binding and the subsequent activation or repression of downstream genes [39]. To determine the direct *functional* targets of AP2XII-8 (i.e., genes that AP2XII-8 has binding affinity to as well as modulations in expression upon binding), we generated scRNA-seq data upon 2 hr AP2XII-8 depletion. Comparing the unperturbed and AP2XII-8 depleted parasites in 3D UMAPs shows a distinct bulge at the G1b-S interface in the KD condition (**Figure 7A**). Under the theoretical assumptions that genes directly controlled by AP2XII-8 reside in the up and down-regulated genes, we performed a DEG analysis, comparing each cell division phase in the AP2XII-8 KD with the corresponding phase in the WT data. This identified a total of 613 DEGs (**Table S4**), of which 95 overlapped with the 246 CUT&RUN identified target genes (**Figure 7B, Figure S7E**). Of the 95 genes in the overlap, referred to as direct functional targets of AP2XII-8 hereinafter, 73 are downregulated (activated by AP2XII-8), 19 are upregulated (repressed by AP2XII-8), while 3 are both down- and up-regulated in different phases (modulated) (**Figure 7C**). Motif searches under the CUT&RUN peaks of activated genes identified 2 motifs (**Figure 7D**): TGCATGCG/A (motif 1) and TATAAGCCG (motif 2). Both motifs were identified in the T_E_1 phase cluster (**Figure 4E**: T_E_1C_E_3C_A_3 which contains only motif 1; T_E_1C_E_2C_A_1 and T_E_1C_E_4C_A_2, which contain only motif 2; T_E_1C_E_3C_A_2 contains both motifs; **Figure S4A**: T_E_1-motif 1 and T_E_1-motif 2 corresponded to the motif 1 and 2, respectively). Thus, these two agnostically detected motifs in the scRNA and scATAC analysis can now be assigned to AP2XII-8.

**Figure 7.**
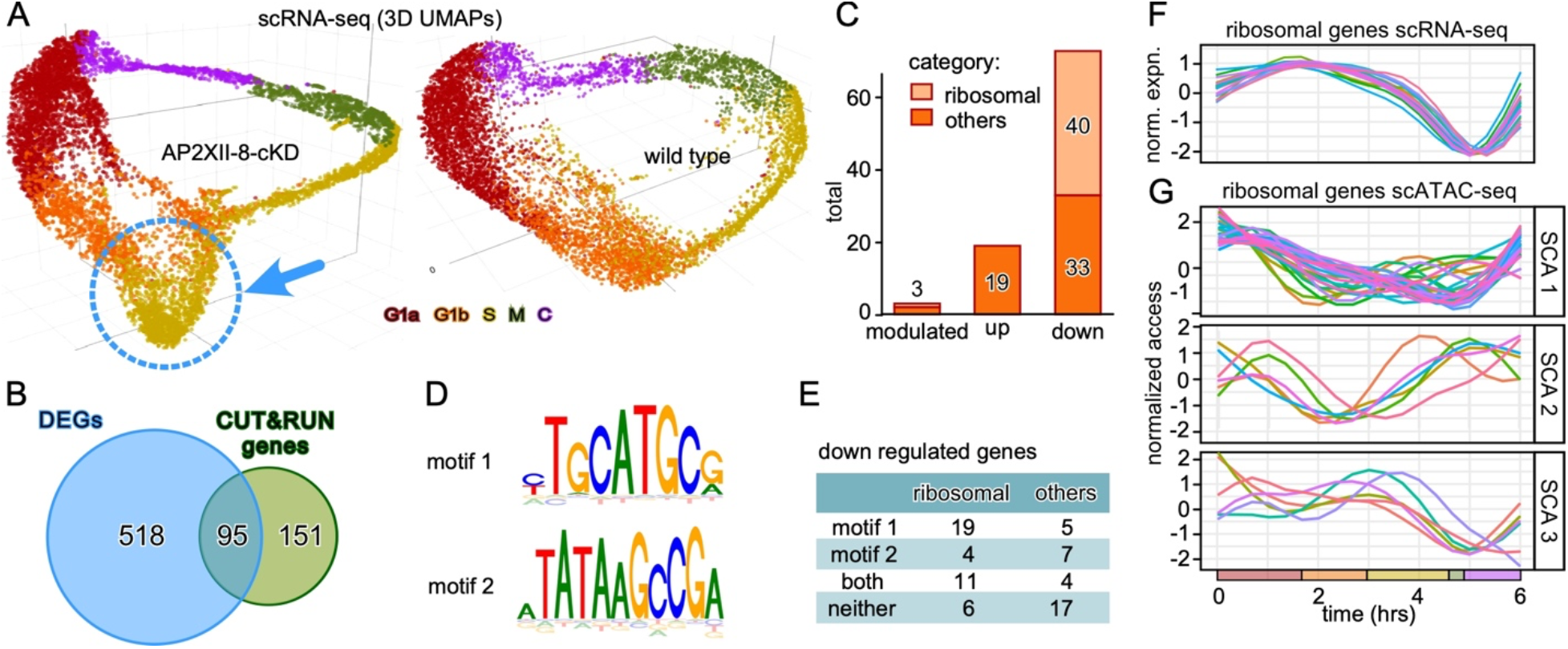
AP2XII-8 perturbed scRNA-seq identifies an AP2XII-8 mediated ribosome regulon. **A.** 3D UMAPs of scRNA-seq data sets collected upon 2 hr KD of AP2XII-8-cKD (left) and wild-type (right, same data as in Figure 2A) parasites. The dotted blue circle marked the ‘bulge cell’ population. **B.** Venn diagram of AP2XII-8-dependent DEGs intersected with the AP2XII-8 CUT&RUN genes. **C.** Differentiation of the 95 genes at the Venn intersection (**B**) by their mode of regulation. ‘modulated’ means either up or down regulated dependent on the cell cycle stage. RP encoding genes are highlighted. **D.** Motif search performed on the 73 AP2XII-8 down regulated genes under the CUT&RUN peaks revealed the presence of two DNA motifs. **E.** Distribution of DNA motif 1 and 2 among the AP2XII-8 down-regulated genes. **F.** Pseudo-time expression profiles of the 40 ribosomal genes under AP2XII-8 control. **G.** Pseudo-time chromatin access profiles of the 40 ribosomal genes under AP2XII-8 control automatically clustered in three sub-clusters (SCA1-3). See also **Figure S7** and **Tables S4**.

### 7. AP2XII-8 controls a ribosome protein regulon

Of the 73 genes activated by AP2XII-8, 40 are ribosomal proteins (RPs). The two motifs engaged by AP2XII-8 were previously mapped in the promoter regions of ribosomal proteins (RPs) [40]: our motif 1 is contained in [T/C] GCATGC[G/A] and motif 2 is a partial match to (TCGGCTTA)TATTCGG (**Figure S7G**). These more extended motifs were named *Toxoplasma* RP promoter elements 2 and 1 (TRP-2 and TRP-1, respectively) [40]. Therefore, we zoomed in on the 40 RPs activated by AP2XII-8 (**Figure 7C**). Compared to the down-regulated non-RPs, the RPs have a nearly two-fold higher incidence of either one or both motifs in their promoters (85% vs 48%: **Figure 7E**). We also checked the specific CUT&RUN enrichment of AP2XII-8 in these RP promoters, which displayed an even stronger mean-intensity than the stringent 246 genes engaged by AP2XII-8 (**Figure S7E** vs **F**). Thus, the specific engagement of these RP promoters by AP2XII-8 is indeed enriched over other RP promoters with these motifs.

The pseudo-time course expression profiles of the AP2XII-8 controlled RP genes are tightly clustered together. They deplete in M-phase and have a broad peak starting at C and around the G1a-G1b transition (**Figure 7F**). The chromatin access profiles of these 40 RPs resolved in 3 clusters each containing considerable variation (sub-cluster accessibility: SCA1-3, **Figure 7G**). Motif distribution did not correlate with the SCA clusters. Therefore, we conclude that AP2XII-8 specifically regulates a sub-set of RPs through a combination of two DNA motifs. This result combined with the notation that our mapped motifs are shorter than the TRP motifs, suggests combinatorial regulation of RPs.

To assess the full extent of AP2XII-8 regulated RPs we performed a motif search for all RPs that have CUT&RUN signal assigned to their promoters regardless of differential expression status (76 out of 137 total RPs). To investigate cooperative binding, we extended the CUT&RUN peaks by 250 nucleotides on each side (determined by examining the tails of CUT&RUN peaks). This identified the two previously reported motifs, matching TRP-1 (TCGGCTTATATTCGG) and TRP-2 ([T/C] GCATGC[G/A]), as well as an additional motif 3 (GAATATAA) not detected under stringent AP2XII-8 CUT&RUN peaks (**Figure S7G**) [40]. Of the 76 RPs with CUT&RUN signal, 72 had at least one of the motifs present with majority of RPs having TRP-2 (**Figure 7H**). Analysis of the distribution of distance between successive motifs shows that all tend to co-occur in relatively close distance to each other, whereas TRP-1 and TRP-2 tend to co-occur close to each other on opposite strands, hinting at possible homo- and/or hetero-dimerization (**Figure S7J-K**; motif 3 was more randomly distributed). Consistent with a role in transcriptional regulation, all three motifs were strongly enriched within 1 kb of the TSS (**Figure S7L**). In conclusion, our analysis maps AP2XII-8 to the control of a subset of RPs through engaging more narrowly defined DNA motifs than the previously mapped TRPs. This is consistent with the presence of additional, combinatorial controllers of RP gene expression.

## Discussion

Combined single cell transcriptome and chromatin landscape analysis of the *T. gondii* lytic cycle facilitated the establishment of highly resolved pseudo-time line at the two major levels of gene regulation. The scRNA and scATAC atlases are accessible through a newly developed web-app that permits data interrogation and visualization in several ways (https://umbibio.math.umb.edu/toxosc/; **Figure 1C**). Chromatin opening within 2 kb of the TSS corresponds very well to the cyclic expression pattern of 90% of the genes (**Figure 2**). The data depth followed by sub-sequent RNA and chromatin level clustering analysis as well as DNA motif mapping permitted resolution of transcriptional regulation of the just-in-time gene expression beyond previous studies [6, 18, 21].

Against the background of known cell cycle checkpoints and the functional modules of apicomplexan cell division (**Figure 1A**) [2, 5, 41, 42], we mapped transition points between gene clusters at the chromatin and transcriptional level (**Figure 3**). The C-G1a transition was consistently present across all levels, even though this is not a formal cell cycle checkpoint. The other checkpoints and cell biological module boundaries were more fluid. Some transitions were only detectable at the chromatin level (e.g. the restriction checkpoint), while others only at the transcriptional level (e.g. spindle assembly checkpoint) (**Figure 3E**). Furthermore, there was one transition that did not align with a cell cycle checkpoint: the T_E_3-T_E_4 transition in the middle of C-phase, which does closely align with the T_A_3-T_A_4 transition. This transition is not clear in the PCA plot, but the scRNA UMAP plot shows two visually distinct cell groups in C-phase: the first is the tail coming out of M-phase, whereas the second half seamlessly associates with the G1a phase (**Figure 2A**). Noting that the assignment of S-phase is based on 2N DNA content [6], these cells are in the middle of the unique intertwined apicomplexan cell division event. At this point, *T. gondii* tachyzoites again execute multiple modules simultaneously: karyokinesis, mother cytoskeleton disassembly, and completion of daughter bud assembly (**Figure 1A**). Indeed, most cortical cytoskeleton assembly genes peak the 5-5.5 hrs mark (BC, alveolins, GAPMs, glideosome, apical cap, sutures). Currently, the genes responsible for karyokinesis and mother cytoskeleton disassembly are largely unknown but must be contained within the roughly 40% hypothetical genes within T_E_3 and/or T_E_4, thereby guiding our future work in revealing this biology. In conclusion, our analysis makes predictions about the level of checkpoint regulation, the modular regulation of transitions, and points at gene clusters containing the genes executing the currently poorly understood biological processes. These findings provide numerous starting points for future dissections of apicomplexan specific biology. The web app permits searching for genes with similar expression and/or chromatin access profiles with variable cut-off similarity settings to guide these research avenues (**Figure 1C**).

The mapping of DNA motifs under the ATAC peaks within 2 kb of the TSS of coordinated gene sets identified several known as well as novel patterns, leading to novel insights on combinatorial regulation of gene expression. The first progressive insight is the identification of three different patterns with a TATA motif (underlined): <u>TATATATAT</u>, <u>TATA</u>AGCC(G), and <u>TATAT</u>GGAT, all of which are in G1 expressed genes. Recently, a TATA box containing motif TCGGCT<u>TATA</u>TTC was mapped by bulk ATAC to 25% of all *T. gondii* genes [21], but it is distinct from the motifs we mapped here. Secondly, we map the TGCATGCA motif to several gene clusters in G1. This motif has been reported before in *T. gondii* [18, 21], and here we mapped it as motif 1 to the AP2XII-8 RP regulon (**Figure 5E**). Furthermore, this regulon is essential for G1 progression and is part of the G1 transcriptional burst.

A combination of two different DNA motifs associated to the same TF has been previously reported for AP2XI-5: GCTAGC is *cis*-activating whereas CAAGAC had no regulatory activity [31]. AP2XII-8 motif 1 is the most abundant in RP genes under its control. We currently do not understand how AP2XII-8 associates with only a subset of the promoters containing either one or both motifs, but we propose a higher-level gene expression control through cooperative binding involving other TFs (e.g., those in AP2 cluster 4, or constitutive TFs) or epigenetic modifiers. Resolving the exact mechanism will be the focus of our future work. Here, we have presented data which strongly support that AP2XII-8 controls an essential RP gene regulon.

TRP-2 is found in genes well-beyond RPs, whereas TRP-1 is more narrowly presents in RP promoters [40]. For example, we see our RP motif 2, in the reverse orientation, in the T_E_1C_E_4C_A_2 cluster, which is associated with small molecule and nucleobase metabolism GO-terms. Indeed, orientation of motifs is a critical factor in how genes are regulated [43]. We derive this is also the case here, and acts as a major factor across the RP promoters. Moreover, we came across another example: we detected AP2IV-4 motif CCCCCCCC in the reverse orientation in a set of T_E_2 genes expressed in S-phase whereas the sense orientation was previously associated with the repression of bradyzoite genes in the acute stage [32] and in addition was also detected in many ribosomal genes [40]. Hence, motif orientation can differentiate activation versus suppression of gene expression.

Aligning with our *T. gondii* data, time course chromatin accessibility and gene expression measurements of synchronized erythrocytic cycle *Plasmodium falciparum* demonstrated that the chromatin opening upstream of transcribed genes corresponds very well with the temporal/developmental changes in transcriptional levels [8]. Indeed, chromatin access displays a slightly wider temporal peak than transcriptional activity in our data. Although at this point no time resolved insights are available, active promotor regions are marked by a complex pattern of histones H3 and H4 methylation and acetylation in both *P. falciparum* [44] and *T. gondii* [20]. Overlaying, integrating and further time resolving these pattens as shown here will advance the understanding of how transcription is organized and meshes at different levels.

In summary, the combination of transcription and chromatin accessibility data synergizes into a deep understanding of the transcriptional network driving the unique *T. gondii* cell division cycle. Furthermore, the new methods and analytical approaches can be can be applied to other biological lytic cycle processes, or to any other comparative cell cycle studies.

## Material and methods

### Parasites and Host Cell Culture

All parasite strains used for this study are listed in the Key resources table. *T. gondii* tachyzoite cultures were maintained in human telomerase reverse transcriptase (hTERT)-immortalized human foreskin fibroblasts (HFFs) as previously described [45]. Immunofluorescence assays (IFAs), were performed using primary HFFs. Selection for stable parasite transfectants was performed under 1 µM pyrimethamine and stable lines were cloned by limiting dilution. Protein knock-downs using the minimal Auxin-Inducible Degron (mAID) system were performed under 500 µM auxin (IAA: 3-indoleacetic acid in 100% ethanol; Sigma-Aldrich).

### Plasmid cloning and parasite strain generation

All primers, synthetic DNA (Twist BioScience), and plasmids used are provided in **Table S6**. All restriction enzyme details are in the Key resources table.

An AP2XII-8 targeted protospacer sequence (P1 primer) was incorporated into *Bsa*I-HF digested pU6-universal plasmid. The resultant plasmid was denoted as pU6-AP2XII-8.

Plasmid mAID-5xTy/DHFR/OsTir1_N was constructed by Gibson assembly (NEBuilder® HiFi DNA Assembly Master Mix; NEB) of the following fragments: 1. mAID CDS PCR amplified (TruFi™ DNA Polymerase; Azura Genomics) from plasmid plinker-AID-Ty-HXGPRT-LoxP using primers P8 and P9; 2. the codon diversified sequence of 5xTy epitope tag including the 3’UTR from DHFR was PCR amplified from pTWIST_5xTY_3’-dhfr using primers P10 and P11; 3. the DHFR promoter was amplified from plasmid DHFR-TetO7-sag4-Ty using primers P12 and P13; 4. the DHFR-TS(m2m3) CDS and T2A skip peptide were amplified from plasmid DHFR_T2A_5xV5 using primers P14 and P15; 5. the OsTir1 CDS was amplified from RHΔ*Ku80*/*Tir1* strain genomic DNA using primers P16 and P17; 6. the HXGPRT 3’-UTR was amplified from a plasmid plinker-AID-Ty-HXGPRT-LoxP using primers P18 and P19.

Subsequently, an *Avr*II restriction enzyme site was introduced in the above plasmid between DHFR-TS and T2A to generate plasmid mAID-5xTy/DHFR/OsTir1, by Gibson assembling the following fragments: 1. plasmid backbone containing mAID-5xTy and the DHFR promoter generated by *Bgl*II, *Aat*II and *Not*I digestion of plasmid mAID-5xTy/DHFR/OsTir1_N; 2. PCR amplified DHFR-TS CDS from plasmid mAID-5xTy/DHFR/OsTir1_N using primers P25 and P26, which introduces the *Avr*II site; 3. PCR amplified fragment containing T2A, OsTir1 and HXGPRT 3’-UTR containing the additional *Avr*II site from plasmid mAID-5xTy/DHFR/OsTir1_N using primers P27 and P28. The resulting plasmid was validated by various restriction enzyme combinations, PCR (primer pairs P22+P23 and P22+P24), Sanger sequencing (primers P6, P21, P21, P22, P23 and P24), and whole plasmid sequencing through Plasmidsaurus.

The AP2XII-8-cKD strain was generated by co-transfecting the RHΔ*Ku80* strain with 50 µg of plasmid pU6-AP2XII-8 and 4 µg of PCR repair cassette amplified from plasmid mAID-5xTy/DHFR/OsTir1 using primers P2 and P3. Parasite clones were validated by diagnostic PCR (**Figure S6**).

### (Immuno-) Fluorescence Microscopy

Intracellular parasites grown overnight in a 6-well plate containing coverslips confluent with HFF cells were fixed with 100% methanol and blocked for 30 min with 0.5% BSA in 1x PBS. The primary antisera (IMC3, 1:2,000; Ty (BB2), 1:500; Centrin (clone 20H5), 1:1,000) and secondary antisera (mouse A594, 1:400; rabbit A488, 1:400) were diluted in blocking solution and applied for 1 hr at RT, followed by three 5 min washes in 1x PBS. DNA staining was performed using 4’,6-diamidino-2-phenylindole (DAPI) in the first wash after the last antibody. Images were acquired with a DeltaVision Ultra microscope and subsequent processing was carried out with FIJI software. For the quantitative analyses for the cell cycle arrest study, at least 100 vacuoles were examined per experiment.

### Western Blot

Intracellularly replicating parasites were harvested through 26G needle passage and filtered (3 µm pore polystyrene filter), washed 3 times in 1X PBS before lysis in resuspension buffer (50 mM Tris-HCl pH 7.3, 150 mM NaCl, 1:50 diluted Protease Inhibitor Cocktail [Sigma-Aldrich, cat log # P8849], 1% SDS). An equivalent of 2×10^7^ parasites whole protein lysate was loaded on a NuPAGE 4-12% Bis-Tris protein gel (Invitrogen, cat log # NP0321BOX) followed by transfer onto a PVDF membrane (Thermo Scientific, cat log # 88520). The blot was blocked in 6% milk in 1X PBS (for α-Ty1 Tag BB2, 1:5,000) or 5% milk 1% BSA (for α-tubulin MAb, 1:2,000) and probed with primary and secondary antibodies (Mouse Immunoglobulins/HRP, 1:10,000) listed in the Key resources table. Blots were washed 3×10 min in PBST (1X PBS with 0.1% Tween 20) following each antibody incubation. Ty signal was developed by SuperSignal West Femto Maximum Sensitivity Substrate (Thermo Scientific, cat log # 34095) whereas tubulin signal was developed by Immobilon Western Chemiluminescent HRP Substrate (Millipore, cat log # WBKLS0100). Chemiluminescent detection was performed on a LI-COR Odyssey M imager. Blots were stripped for re-probing using stripping buffer (62.5 mM Tris-HCl pH 6.8, 2% SDS, and 0.8% (v/v) β-mercaptoethanol) at 50°C for 20 min followed by 3×10 min PBST washes.

### scRNA-seq

RHΔ*Ku80*::ptub-YFP_2_/sagCAT (WT) or AP2XII-8-cKD parasites were grown overnight in hTERT cells and harvested by 26G needle passage and 3 µm pore polystyrene filtration. Parasite concentrations were adjusted to 2.5×10^6^/ml and 4 ml (10^7^ tachyzoites) were inoculated on T25 flasks confluent with HFFs. Two T25 flasks were used per time point (27, 30, and 33 hpi for WT; 27, 30 hpi for AP2XII-8-cKD; IAA was applied for 2 hrs to AP2XII-8-cKD parasites prior harvest) and intracellular tachyzoites harvested by washing monolayers 3X with ice-cold 1X PBS, scraping the parasites-contained HFFs in ice-cold Ed1 media, and releasing parasites by 26G needle passage. For each time point 5×10^6^ tachyzoites were each pelleted at 1,000x*g*, 4°C for 15 min and resuspended in 1 ml of ice-cold resuspension buffer (1X Hank’s Balanced Salt Solution (Thomas Scientific, Cat#SH30268.02), 0.04% BSA (MilliporeSigma-Omnipour, Cat#29-301-00GM)). Parasites were pelleted at 1,000x*g*, 4 °C for 15 min and resuspended in 60 µl of ice-cold resuspension buffer. Recounted parasites were pooled across the time points at 10^6^/ml for at an end volume of 150 µl.

scRNA-seq library preparation followed the Chromium Next GEM Single Cell 3’ Reagent Kits v3.1 protocol (10x Genomics, CG000204 Rev D; WT) or Chromium Next GEM Single Cell 3ʹ Reagent Kits v3.1 (Dual Index) (10x Genomics, CG000315 Rev D; AP2XII-8-cKD), aiming for 10,000 cell recovery. The quality and concentration of the generated scRNA-seq library was assessed using an Agilent 2200 TapeStation. The WT library was sequenced on an Illumina NextSeq 550 High-Output Kit (150 cycles) flow cell at the Tufts University Core Facility (TUCF) with the following sequencing parameters: Read 1 (28 cycles), i7 Index (8 cycles), i5 Index (0 cycles), and Read 2 (91 cycles). The AP2XII-8-cKD library was sequenced on an Illumina NextSeq 550 Mid-Output Kit (150 cycles) flow cell at TUCF with the following sequencing parameters: Read 1 (28 cycles), i7 Index (10 cycles), i5 Index (10 cycles), and Read 2 (90 cycles).

### scATAC-seq

RHΔ*Ku80*::ptub-YFP_2_/sagCAT inocula were prepared as for scRNA-seq. 5×10^6^ mechanically released tachyzoites per time point were each pelleted at 1,000x*g*, 4°C for 15 min. Without disturbance, the pellet was washed with 100 µl ice-cold 1X PBS followed by 1,000x*g*, 4°C for 15 min. Cells were lysed using 100 µl of ice-cold ATAC lysis buffer (Active Motif), followed immediately by addition of 1 ml pre-chilled wash buffer (10 mM Tris-HCl pH 7.4, 10 mM NaCl, 3 mM MgCl_2_, 1% BSA, 0.1% Tween-20). Nuclei were pelleted at 500x*g*, 4°C for 5 min and resuspended in 190 µl ice-cold resuspension buffer (as for scRNA-seq). Each sample was resuspended to 4×10^6^/ml and 100 µl of each nuclei sample was pooled.

The scATAC-seq library was produced following the Chromium Next GEM Single Cell ATAC Reagent Kits v1.1 protocol (10x Genomics, CG000209 Rev G) aiming for 10,000 nuclei recovery. The quality and concentration of the generated scATAC-seq library was assessed on an Agilent 2100 Bioanalyzer and sequenced on a NovaSeq 6000 platform at the Bauer Core Facility (Harvard University) using the following run parameters: Read 1N (50 cycles), i7 Index (8 cycles), i5 Index (16 cycles), and Read 2N (50 cycles).

### CUT&RUN

Intracellularly replicating AP2XII-cKD or RHΔ*Ku80* parasites were harvested 30 hrs post-inoculation by 26G needle passage and pelleted by centrifugation at 1,000*x*g, 4°C for 15 min (same spin condition for subsequent washes). Parasites were washed following the ChIC/CUT&RUN Kit Version 3 (EpiCypher, Inc, Catalog No. 14-1048) user manual before the binding to Ty1 BB2 MAb, Ty1 Diagenode MAb Antibody, Mouse IgG1 Isotype Control (antibody details in the Key resources table and **Figure S7A-D**). Subsequent chromatin digestions and CUT&RUN DNA generations were performed following the user manual (EpiCypher), with 5 ng CUT&RUN-enriched DNA carried over to the library preparation step.

The CUT&RUN libraries were produced utilizing the CUT&RUN Library Prep Kit (EpiCypher, Inc, Catalog No. 14-1001) User Manual Version 1.0, with modifications pointed out in the manual tailored specifically to the transcription factor applications. The qualities and concentrations of the generated CUT&RUN libraries were assessed using an Agilent 4150 TapeStation. The libraries were multiplexed following the Primer Selection Guide (EpiCypher, Inc) and sequenced on the Illumina MiSeq Standard V3 150 cycles (75 bp paired-end) flow cells at the Tufts University Core Facility (TUCF).

### RNA- and ATAC-seq read alignment

Reference genome and gene annotations for *T. gondii* (ME49 strain, release 59) were downloaded from https://toxodb.org/ [46]. Custom references were assembled using the 10X genomics pipelines (cellranger-7.0.1 and cellranger-atac-2.1.0) [47] using the cellranger mkref command. Subsequently, the raw RNA-seq and ATAC-seq reads were mapped against the reference genome using cellranger count and cellranger-atac count commands respectively with the default parameters.

### scRNA-seq data processing

Downstream data analysis was performed in R (version 4.2.2). Gene expression count data was processed using the Seurat R package (version 4.3.0) [23]. Lowly expressed genes and cells with a few number of detected reads were filtered from downstream analysis using Seurat function CreateSeuratObject with parameters min.cells = 5, and min.features = 100. Expression data was normalized and scaled using NormalizeData with parameter normalization.method = “LogNormalize”, FindVariableFeatures with nfeatures = 6000, and ScaleData. Dimensionality reduction was performed using Principal Component Analysis (PCA) and Uniform Manifold Approximation (UMAP) as implemented in Seurat using runPCA() and runUMAP() with parameter dims = 1:10. Clustering analysis was carried out using KNN graph based technique using FindNeighbors and FindClusters functions with parameters dims = 1:10, reduction = “pca”. The data set was down sampled to include 8000 cells. Cell labels were inferred using the published scRNA-seq data [6] with Seurat functions FindTransferAnchors() and TransferData().

### scATAC-seq data processing

The chromatin accessibility peak-by-cell matrix data was processed in R using the Seurat R package (version 4.3.0) [23]. Cells and peaks with low counts were filtered out using Seurat function CreateChromatinAssay with parameters min.cells = 5 and min.features = 100. Significantly enriched peaks were selected for further downstream analysis (peak_region_fragments > 200, peak_region_fragments < 6000, pct_reads_in_peaks > 40), nucleosome_signal < 4 and TSS.enrichment > 2). Data was normalized with Seurat function RunTFIDF. Dimensionality reduction was performed using singular value decomposition (SVD) and Uniform Manifold Approximation (UMAP) as implemented in Seurat functions RunSVD with default parameters and RunUMAP with parameters reduction = ‘lsi’ and dims = 1:30. Clustering analysis was performed with functions FindNeighbors and FindClusters with parameters reduction = ‘lsi’, dims = 1:30 and algorithm = ‘SLM algorithm’. Gene activity matrix was then generated from the chromatin assay using GeneActivity function extending the regions to 600 and 200 bp up- and down-stream of the TSS. The gene activity was utilized to create a Seurat object followed by normalization and clustering analysis similar to the scRNA-seq data. The scRNA-seq and scATAC-seq data were integrated using Seurat functions SelectIntegrationFeatures, FindIntegrationAnchors and IntegrateData with scRNA-seq data used as reference for integration.

### Peak gene assignment

All peaks detected by cellranger 10x pipeline (5444 total peaks) and gene annotation file were used to assign peaks to genes using bedtools [48]. Peaks located entirely within the gene/coding region (exon-2 to exon-n) were filtered out from further analysis (427 total). The remining peaks were assigned to the nearest downstream gene with distance cut-off < 3000 bp. Multiple peaks assigned to the same gene were merged. In total 5322 genes were uniquely assigned to a peak.

### Pseudo-time analysis

Pseudo-time analysis was performed as previously described [22]. Briefly, an elliptic curve was fitted to the first two principal components in each scRNA- and scATAC-seq data using Ellipsefit function from MyEllipsefit R package (https://github.com/MarkusLoew/MyEllipsefit). The elliptic curve was used as a prior to fit a closed principal curve to each data set using principal_curve function from the R package princurve with the parameter smoother = “periodic_lowess” [49]. Cells were orthogonally projected on to the principal curve and were ordered according to arc-length distance from the beginning of the curve. The transferred cell cycle phases were used to adjust the start time of the trajectory to match the start of G1a phase. A piecewise linear scaling was performed to scale each phase to match the actual length as previously reported [4].

### Marker analysis

Differential expression analysis (DEA) was performed using FindAllMarkers function from the Seurat R package with parameters only.pos = TRUE. This analysis was carried out to identify DEGs of each inferred cell cycle phase. Fold change cutoff > 1.5 and adjusted p-value < 0.05 were used to determine significant DEGs. The same function and cutoffs were used to identify DEGs of each detected transition group.

### Fitting gene curves

The expression and accessibility of genes along the reconstructed pseudo-time were estimated by fitting a smoothing spline to expression and accessibility profile of each gene with smooth.spline function and smoothing parameter *λ* = 0.1 and weight = 1/3 for 0 expression values. The fitted splines were sampled at regular intervals. Timing of peak expression and accessibility per gene were calculated by examining the local maxima of the fitted curves.

### Curve cross correlation analysis

Reconstructed time course expression and accessibility were used to measure the similarity (correlation) of the two curves in corresponding genes using ccf function in R [50].

### Transition points along the cell cycle

A smoothing spline was fitted to the calculated peak expression and peak accessibility of cell cycle regulated genes with the smooth.spline function and smoothing parameter *λ* = 0.005. The smoothing parameter was set by trial and error. The transition points in the peak expression and peak accessibility curves were identified by examining the inflection points of the fitted curves. The detected points of inflection were used to calibrate transition time during cell cycle.

### Time course clustering

Time course clustering of gene expression curves was performed as previously described [22]. Briefly, gene expression and accessibility curves were individually utilized to build *n* × *N* time course matrices, with rows representing genes and columns representing expression/accessibility time points. Each data was z-score scaled and a Dynamic Time-Warping (DTW) clustering algorithm was used to measure the similarities between the time series. In this analysis we used the function tsclust from dtwclust R package (https://github.com/asardaes/dtwclust) with the following parameters: type = hierarchical, control = hierarchical_control(method = “complete”), args = tsclust_args(dist = list(window.size = 4L). The number of clusters was set by empirically trial and error.

### RNA Velocity calculations

For each data set, loom files were generated using velocyto command line tool [34]. Loom files were processed in python using the scanpy library [51] and spliced/unspliced counts were estimated. The scvelo python package [35] was used to calculate velocity lengths using the dynamic model.

### CUT&RUN data analysis

Paired-end CUT&RUN reads were mapped with BWA-mem (version 0.7.17) [52] against the *T. gondii* ME49 reference genome. Peaks were called relative to the negative control with MACS2 [53] using the following command and parameters: macs2 callpeak -p 0.05 -m 2 50 -t AP2XII-8.sorted.bam -c RH_Negative.sorted.bam --nomodel -f BAM --extsize 120.

### Gene set enrichment

Gene Ontology Enrichment Analysis (GOEA) was performed using available GO on ToxoDB.org (version 64) and significant GO terms (Benjamini < 0.1) were determined.

### Motif search

Markers of each transition group were identified. Motif search was performed under the ATAC peak of marker genes using BAMM (https://bammmotif.soedinglab.org/)

## Data and code availability

An interactive web-application for visualization and exploration of our data set can be found here: https://umbibio.math.umb.edu/toxosc/. The analysis R code is available on GitHub: https://github.com/umbibio/scToxoplasmaCDC (DOI: <u>10.5281/zenodo.8219739</u>). scRNA-seq and CUT&RUN data (fastq) have been deposited to the Sequence Read Archive (SRA) under the accession number SUB13707798.

## Acknowledgements

We would like to thank VEuPathDB for the readily accessible *T. gondii* genome assemblies and their functional annotations. We would also like to thank David Roos for helpful suggestion regarding the analysis, particularly relating to grouping genes based on expression/accessibility profile similarities.

This study was supported by the National Institute of Health grants AI150090 (KZ and MJG), and AI167570 (KZ, MTD and MJG). We would like to acknowledge the 1S10OD032203-01 (Tufts University Core Facility Genomics Core) for the support of the included NGS analysis. The funders had no role in study design, data collection and analysis, decision to publish, or preparation of the manuscript.

## Author Contributions

KZ, MTD and MJG conceived the approach, KZ and MJG co-wrote the manuscript. JL performed all wet-lab *T. gondii* experiments, and established scRNA-seq and scATAC-seq in collaboration with CDK. YR, AA, and KZ performed data analysis. DD and KZ developed periodic spline fits. All authors edited and proofread the final manuscript.

## Supplementary material

### Supplementary Tables

**Table S1.** Quality control. Tabs 1 and 2 contain the sequencing and alignment metrics of scRNA- and scATAC-seq data, respectively. Tab 3 summarizes the mapping rates of CUT&RUN samples to the reference genome.

**Table S2.** Differential expression. Tabs 1 & 2 provides list of DEGs DEGs within each canonical cell cycle phases and inferred transition points (FC > 1.5 and adjusted p-value < 0.05).

**Table S3.** Gene clusters. Tab 1 contains list of genes in each RNA and ATAC sub clusters.

**Table S4.** AP2XII-8 target genes. Tab 1 contains the list of genes targeted by AP2XII-8 detected by analysis of CUT&RUN data sets. This also contains the list of DEGs upon AP2XII-8 KD. Motifs enriched under the CUT&RUN peaks of these genes are shown.

**Table S5.** ATAC peaks, Tab 1 contains list of peaks that were assigned to the nearest downstream gene with the distance cut-off < 3,000 bp.

**Table S6.** All oligonucleotides and recombinant DNA used in this study. Sequences are shown in 5’-to 3’-orientation. ‘Lab ID’ refers to primer log in the Gubbels lab.

**Table S7.** Quantification of cell cycle arrest stages upon AP2XII-8 KD.

## Supplementary Information (SI) Figures

**Figure S1.**
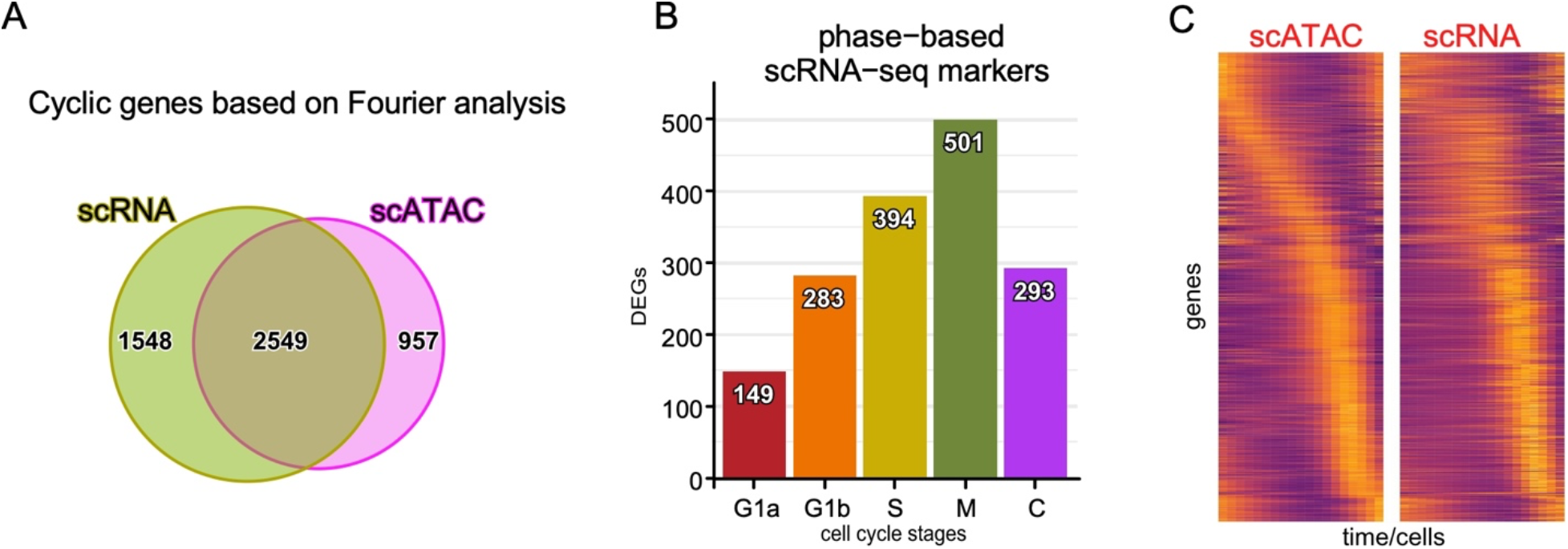
Uncovering differentially expressed cyclic genes, Related to Figures 2 and 3. **A.** Fourier-based analysis identified 4,097 and 3,506 cyclic genes according to their expression and accessibility profiles, respectively. Ven diagram display 2,549 genes which were cyclic in both expression and accessibility curves. **B.** DEGs within each inferred cell cycle phase (FC > 1.5, adj-p-value < 0.05). **C.** Pseudo-time heatmaps of single cell transcriptome (right) and chromatin access (left) for the 1,238 genes displaying cyclic expression profiles. Top to bottom organized by timing of peak detection in scATAC-seq data.

**Figure S2.**
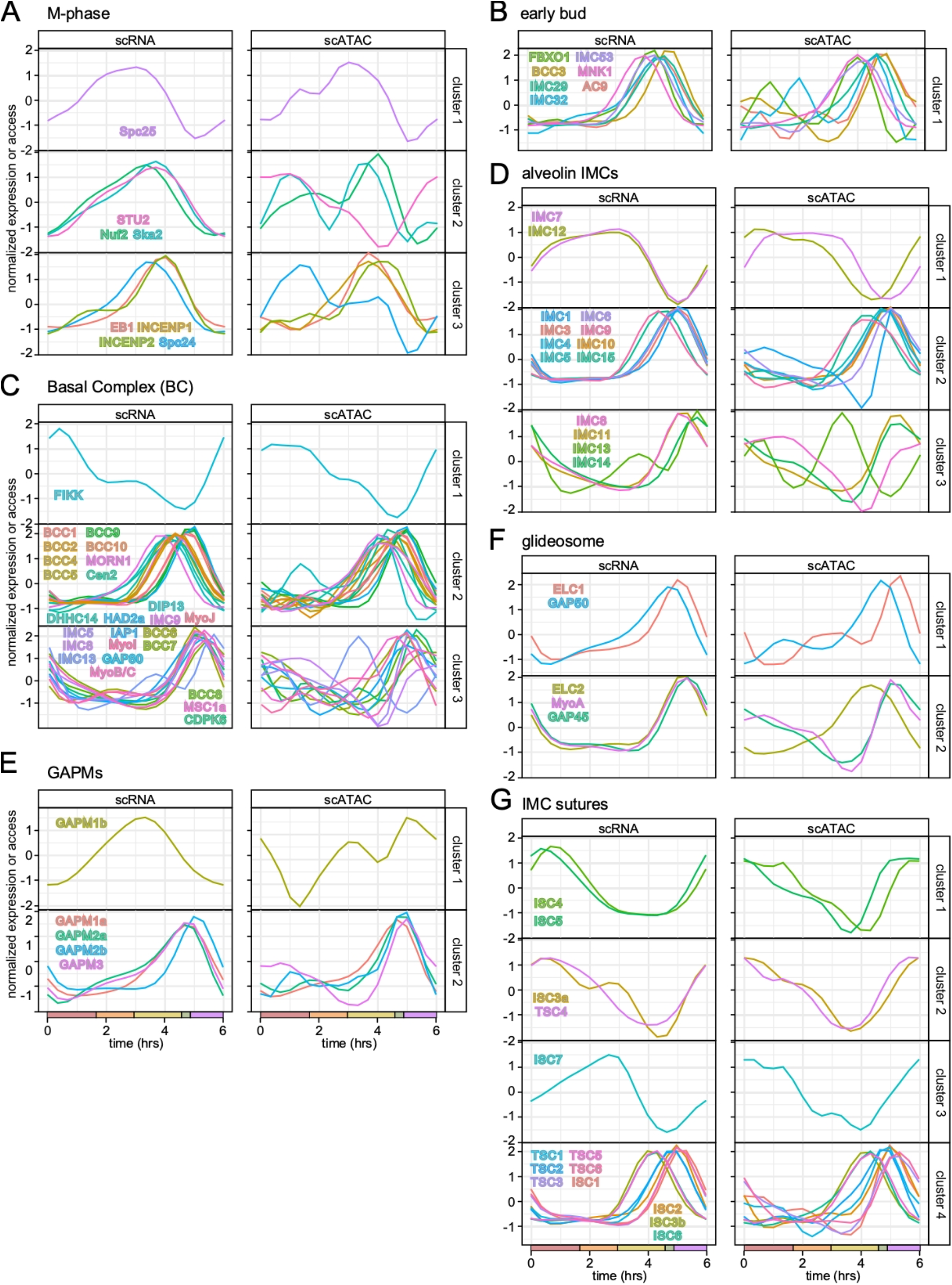
Representative chromatin accessibility and expression patterns of functional groups, Related to Figure 4. Profiles are automatically clustered based on their expression patterns (left) and shown together with their corresponding chromatin access profiles plotted on the right. **A.** M-phase genes **B.** Early bud genes **C.** Basal complex (BC) genes. BC composition and function changes through the cell division cycle [26], which is reflected in the expression patterns. **D.** Alveolin domain containing IMC proteins. Alveolin gene expression patterns match their experimentally validated sequential assembly into the IMC [28]. **E.** Gliding-associated membrane proteins (GAPM) genes connect the IMC to the subpellicular microtubules [56]. **F.** Glideosome associated genes. **G.** IMC suture genes. The IMC sutures are found inbetween the alveolar vesicles making up the cortical membrane skeleton [29, 30].

**Figure S3.**
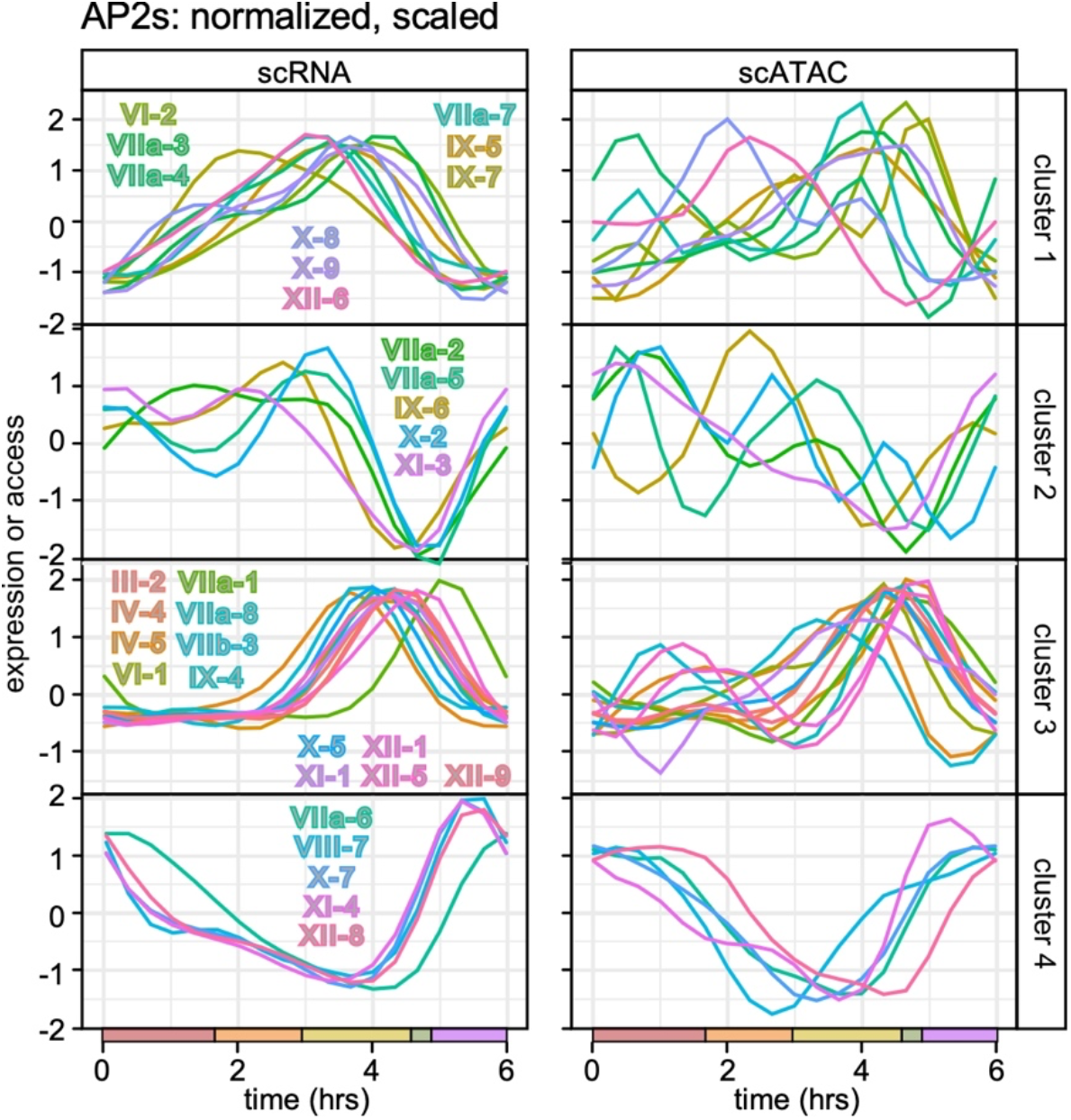
Chromatin accessibility and expression patterns of the 32 cyclic AP2 TFs, Related to Figure 4. Profiles are automatically clustered (clusters 1-4) based on their expression patterns (left), and shown together with their corresponding chromatin access profiles plotted on the right.

**Figure S4.**
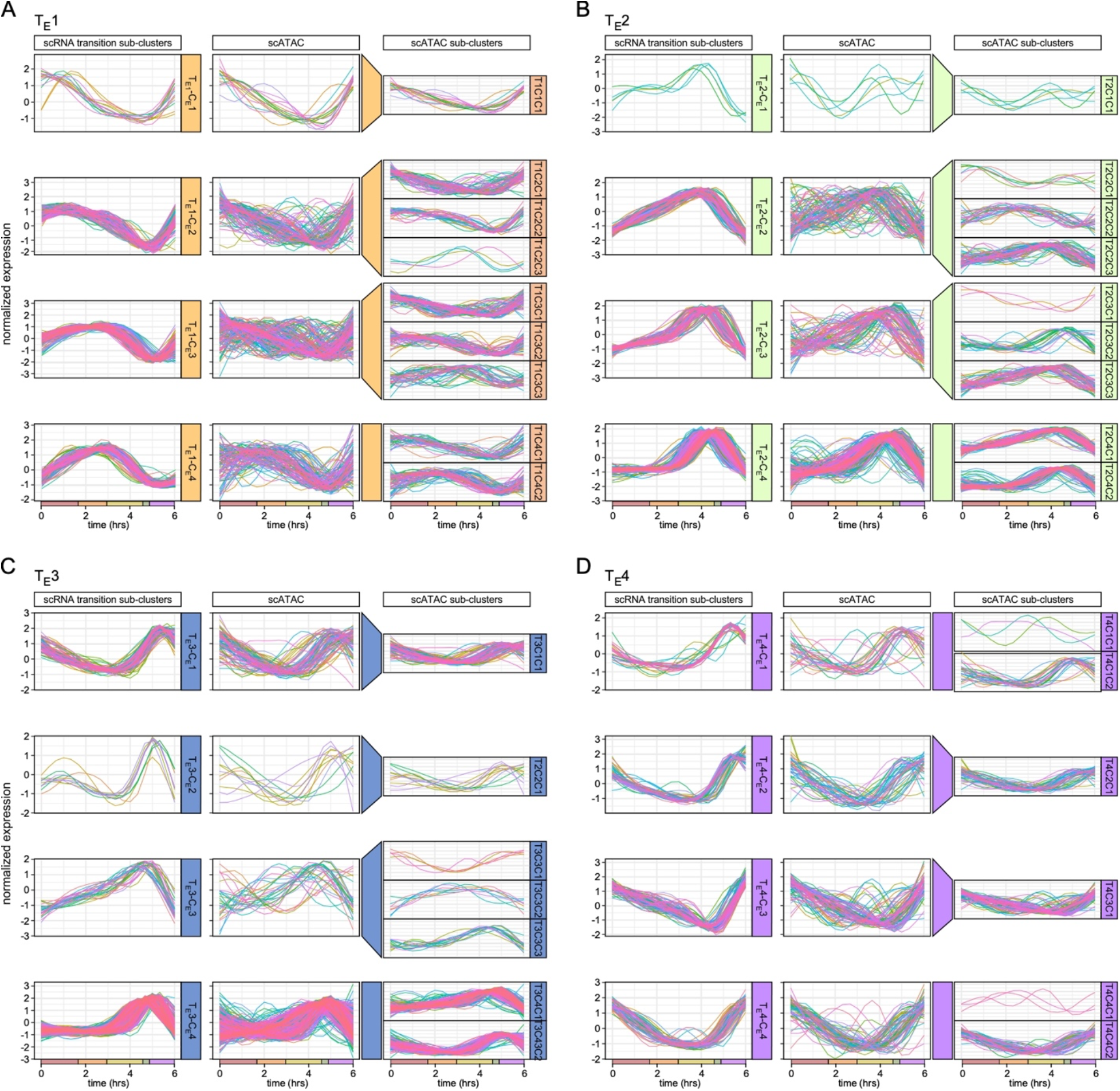
Clustering analyses of genes in transition expression clusters T_E_1-4 based on their expression and chromatin access profiles, Related to Figure 4. DEGs in each expression-derived transition cluster (Figure 3F), T_E_1-4 **(A-D)**, were separated into four sub-clusters (C1-4) based on the shapes of their scRNA-seq time resolved expression profiles, shown on the left. The corresponding scATAC-plots for the genes in each scRNA transition sub-cluster are shown in the middle, which were subsequently automatically sub-clustered on the far right.

**Figure S5.**
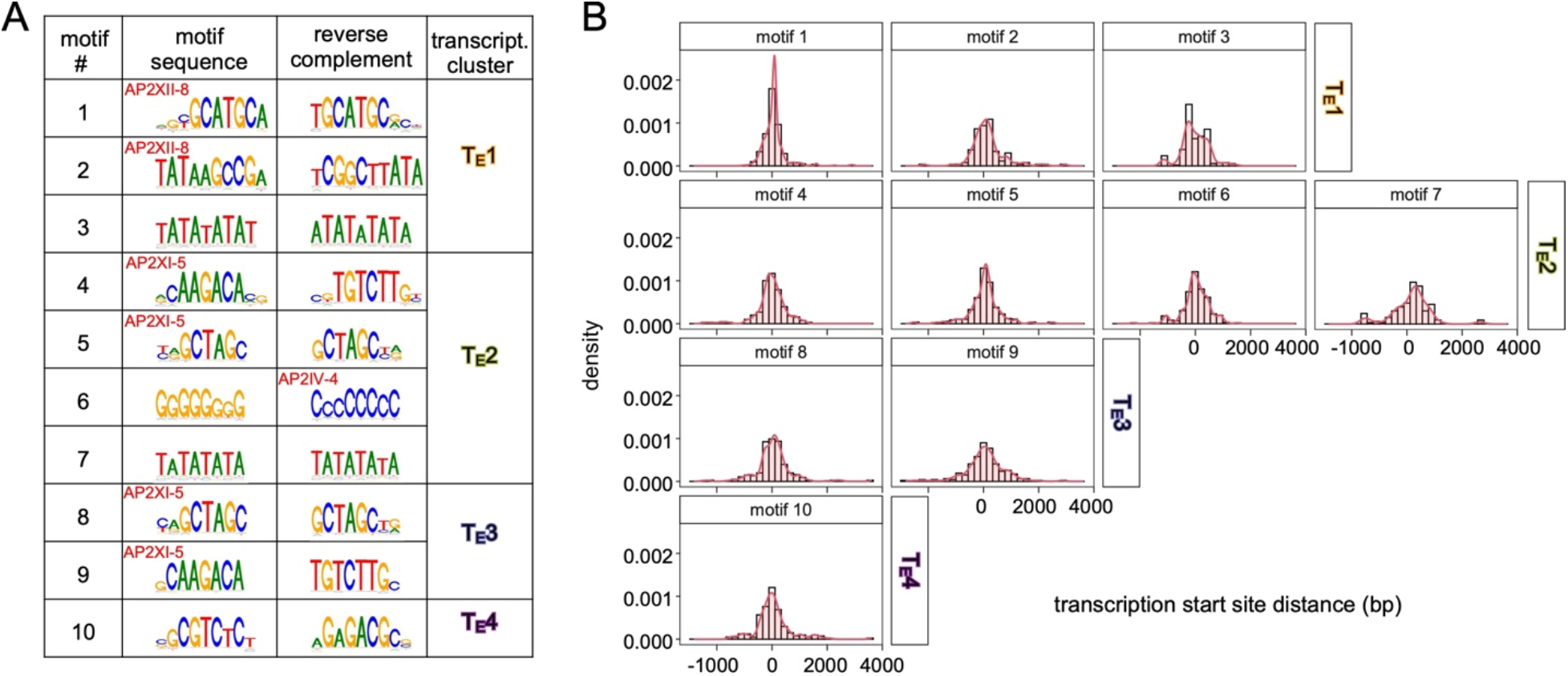
Motif mapping for co-expressed/co-accessible genes in expression transition clusters T_E_1-4, Related to Figure 4. **A.** Ten motifs were identified for the DEGs in each expression-derived transition cluster (Figure 3F) under the ATAC-peaks. These included several previously mapped motifs recognized by specific AP2 TFs as indicated. **B.** Each motif was mapped on the genes where it was found; the transcription start site (TSS) is set at 0 and plots show up to 4000 bp up- or down-stream as relevant for each factor.

**Figure S6.**
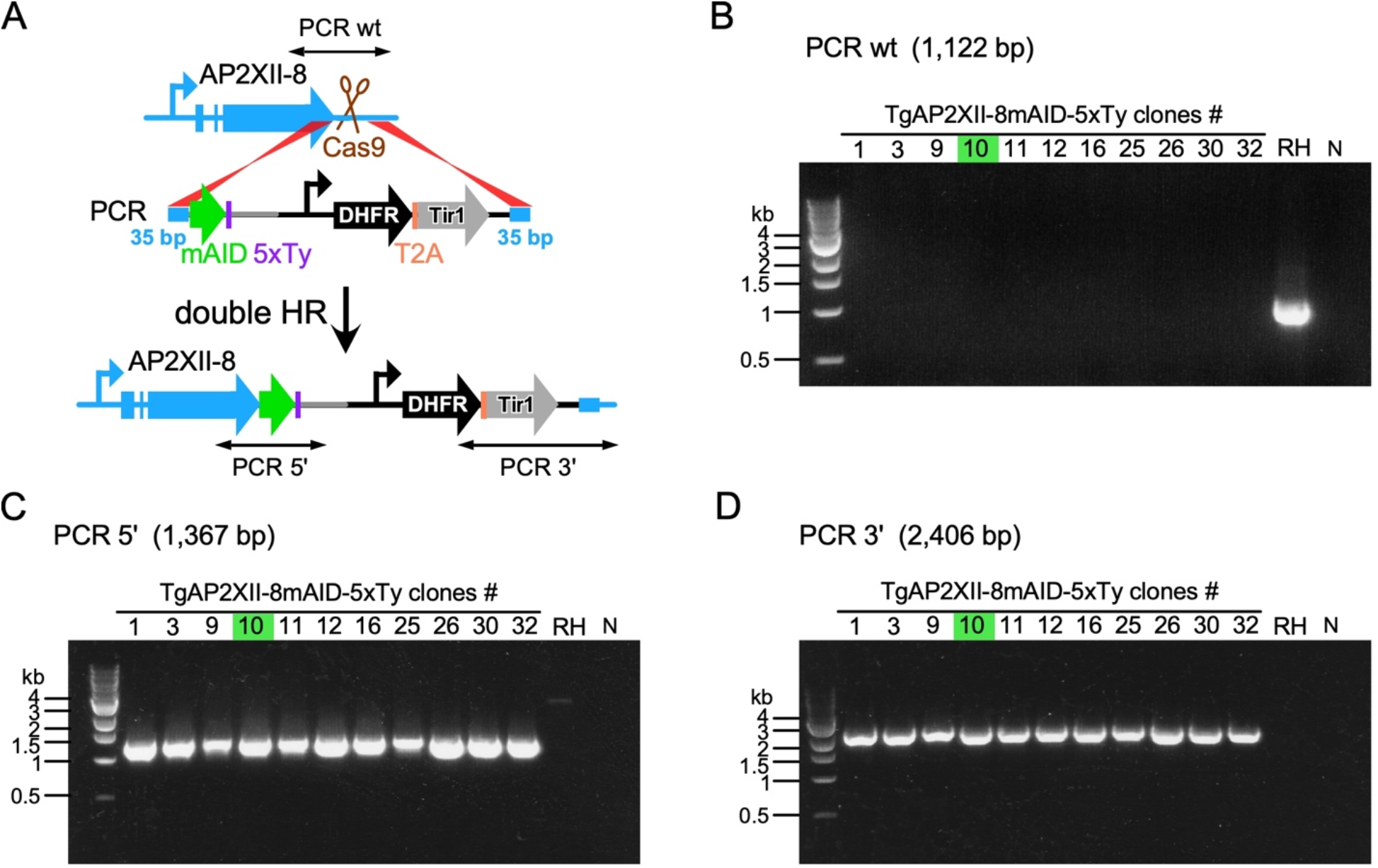
AP2XII-8-mAID-5xTy/Tir1 generation and PCR validation, Related to Figure 5. **A.** Schematic representation of the one step strategy that simultaneously tags AP2XII-8 at the C-terminus with mAID-5xTy and introduces the DHFR selectable marker fused to the Tir1 open reading frame through a self-cleaving Thoseaasigna virus 2A (T2A) peptide. Double homologous recombination (HR) is facilitated by 35 bp homologous flanks and a CRISPR/Cas9 generated dsDNA break. **B-D.** Diagnostic PCR reactions as indicated on single parasite clones (numbered), RHΔ*Ku80* (RH) parent strain, and a no DNA negative (N) control. Expected lengths as indicated. Primer pair localizations are marked in panel **A** using the following primer pairs: PCR wt: primers P4 and P5; PCR 5’: primers P4 and P7; PCR 3’: primers P5 and P6 (see Key resources table for primer sequences). Clone #10 (green) was used in all further experiments.

**Figure S7.**
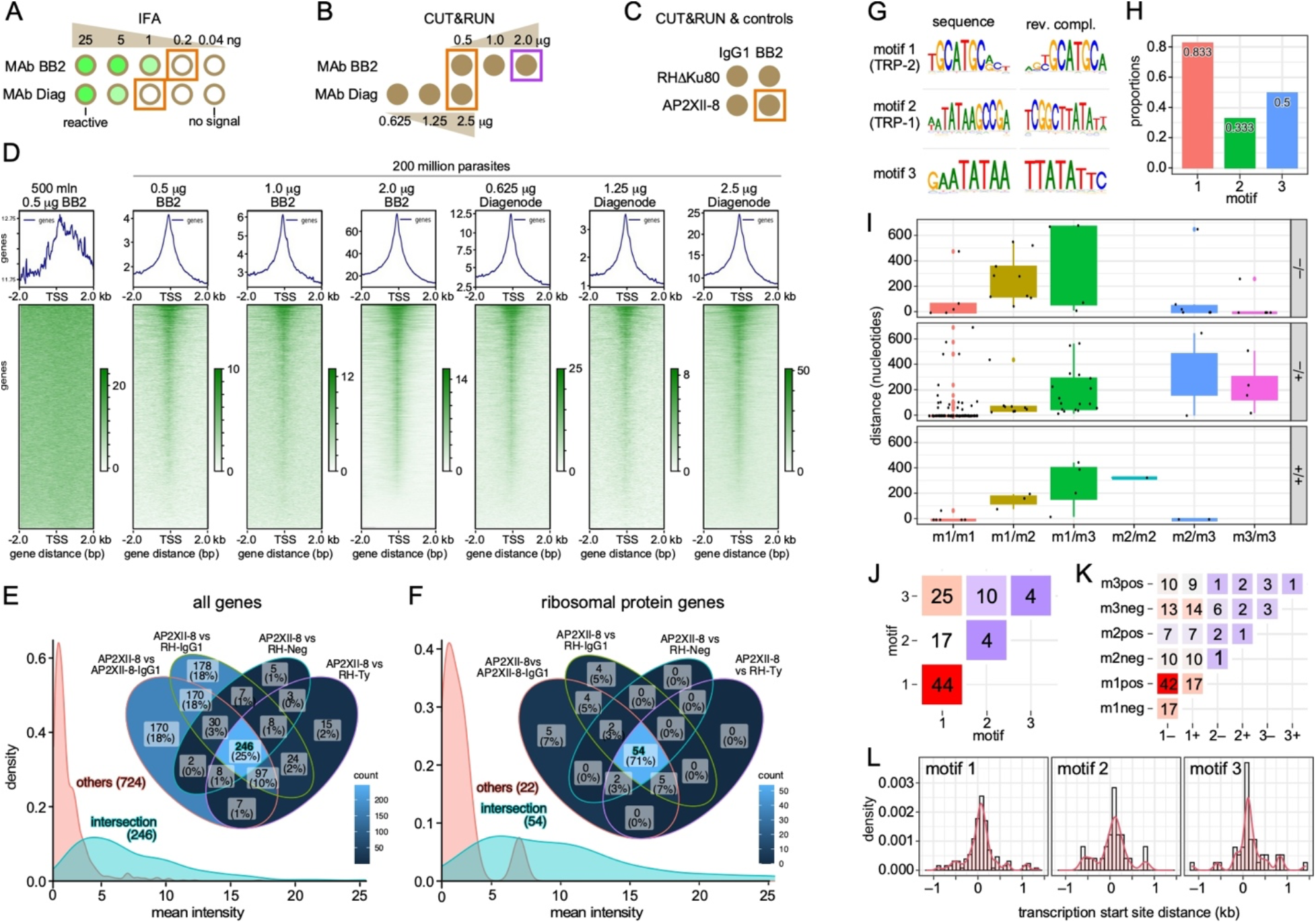
CUT&RUN optimization, controls, and analysis of motif co-occurrence under the CUT&RUN peaks of RPs, Related to Figures 6 and 7. **A.** Two different Ty-epitope specific monoclonal antibodies (MAb) were first compared for affinity by an IFA antibody dilution series on AP2XII-8-mAID-5xTy expressing parasites. The orange boxed dilutions defined the first value below the detection limit. BB2 is the standard antiserum used to detect Ty in IFA and western blots, whereas the ‘Diag’ antibody is marketed for Ty based ChIP experiments, which is in experimental principle comparable to CUT&RUN. **B.** The two MAbs were used in two different dilution series above and beyond the value recommended by the manufacturer. The orange boxed region is functionally equivalent as derived from the dilution series in panel **A**. The purple boxed value was selected for subsequent experiments. **C.** Experimental design of the definitive CUT&RUN experiments, including the negative controls. 2.0 μg BB2 MAb or IgG1 isotype control was used on 200 million parent (RH1′Ku80) or AP2XII-cKD parasites harvested from asynchronously replicating cultures. **D.** Heatmap of the CUT&RUN reads assembled around the TSS for the various controls and experimental samples under the conditions as indicated. **E, F.** Venn diagrams of the unique genes mapped to each CUT&RUN sample comparison as indicated. Panel **E** comprises all genes (970), while panel **F** comprises of only the ribosomal genes (76). The density profiles at the bottom display the mean intensity profiles of the loosely controlled genes (only genes found in all controls excluded) vs the most tightly (middle intersection in the Venn diagram; any gene detected in any of the negative controls excluded) selected CUT&RUN genes. **G.** Motifs identified under the CUT&RUN peaks of all RPs with signal over any of the negative sets. Motif-1 and Motif-2 are previously reported as *T. gondii* TRP-2 and TRP-1, respectively [40]. **H.** Proportion of RPs having the indicated motif. **I.** Distance of successive motifs identified under the same CUT&RUN peaks. Facets indicate the strand on which the successive motifs were identified. **J.** Heatmap shows motif co-occurrence. Motif 1 co-occurs highly with itself and motif 3. **K.** Heatmap shows motif co-occurrence as in **J**, but considering strand of the motif. **L.** Motif distance to the TSS of the ribosomal genes.

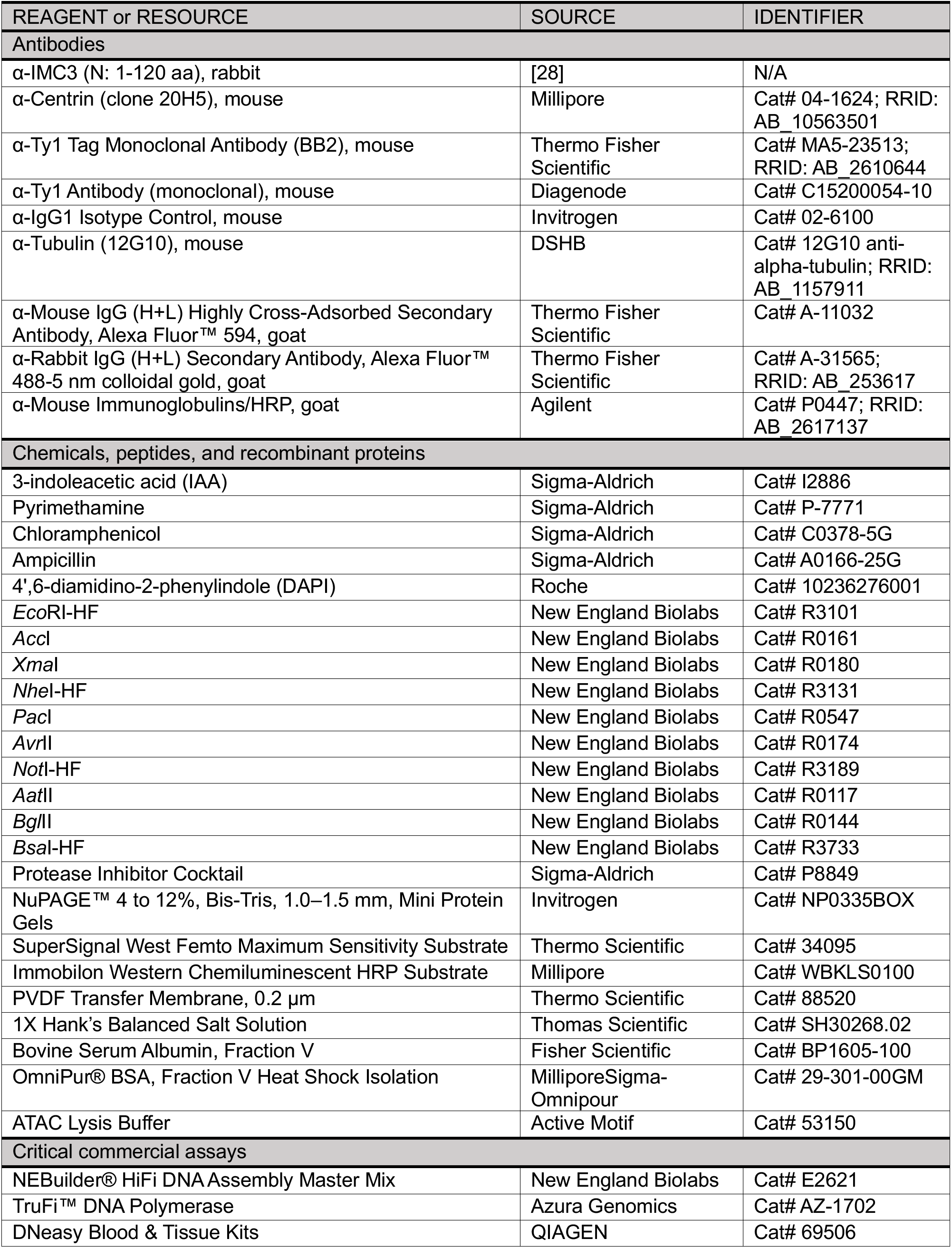
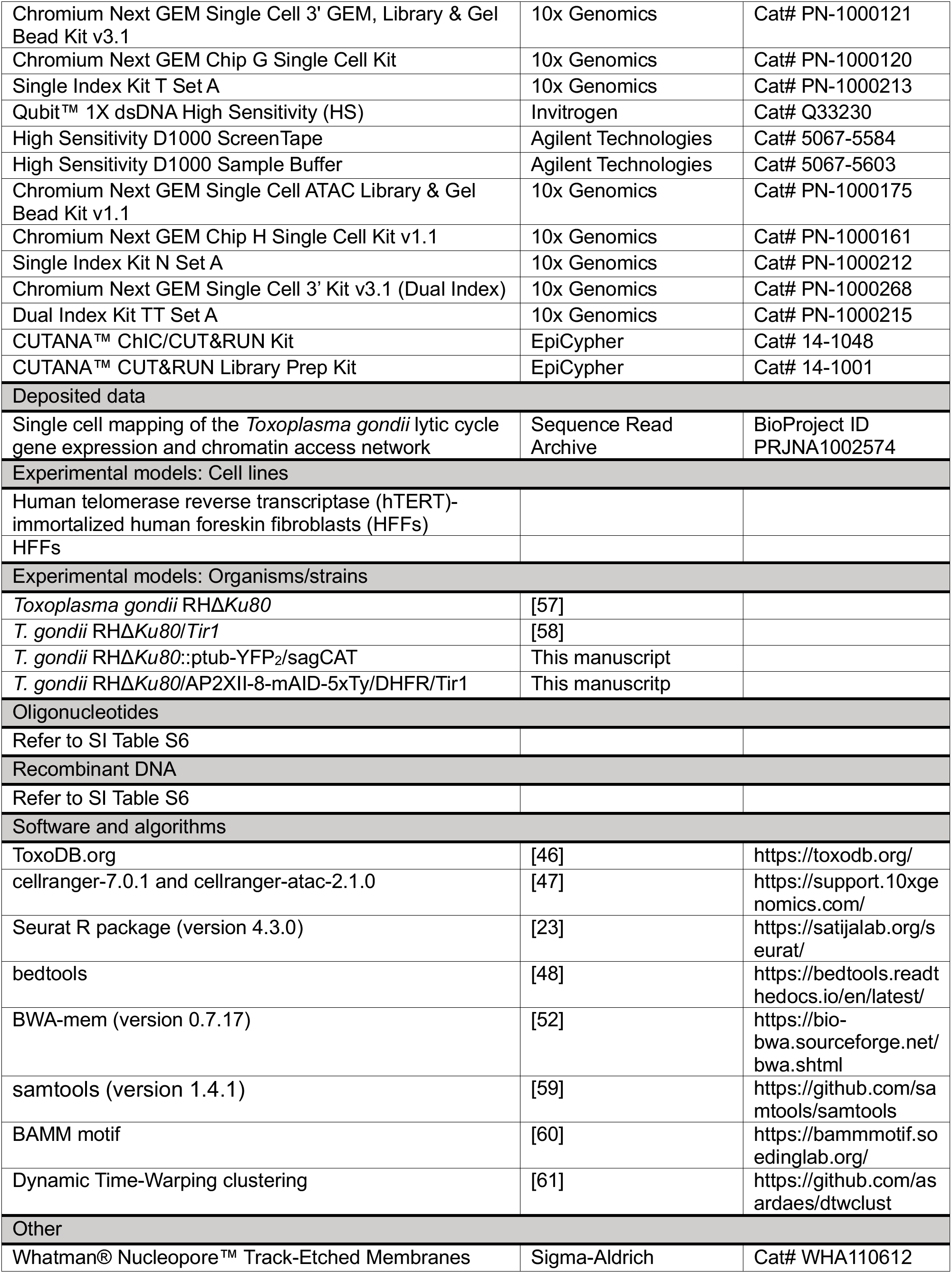
Key resources table.

